# Deep learning design and *in vivo* validation of Müller glia-specific *cis*-regulatory elements

**DOI:** 10.64898/2026.07.15.738785

**Authors:** Oriol Fornes, Kyung Duk Koh, Rebecca M. Marton, Jiapei Chen, Carlo De Donno, Kevin A. Marroquin, Gyeong Jin Kang, Avantika Lal, Sören Müller, Gokcen Eraslan, Eliah R. Shamir, Pratiksha I. Thakore, Heinrich Jasper, David Garfield

## Abstract

Effective recombinant adeno-associated virus gene therapies require promoters that are compact and cell-type-specific. Ideally, promoters should also exhibit functional conservation when tested in model organisms to ensure that preclinical findings translate reliably to human patients. Here, we introduce a deep learning framework for designing *cis*-regulatory elements (CREs) meeting these criteria, applied to retinal Müller glia (MG). Using single-cell chromatin accessibility data from human and mouse retinas, we trained species-specific models to predict cell-type accessibility, and designed compact CREs using two complementary strategies. *In silico* validation predicted that the designed CREs exhibit high MG-specific accessibility (on-target) in both species with minimal off-target accessibility across hundreds of human cell types and tissues. Mechanistic analysis revealed that the predicted MG accessibility is driven by the creation of LHX2 motifs. *In vivo* validation confirmed that the designed CREs successfully restrict reporter expression to MG in the murine retina. Our deep learning framework is highly generalizable and enables the rapid design of compact, on-target, and species-conserved CREs for precision gene therapy.

## Introduction

Recombinant adeno-associated virus (rAAV) vectors have emerged as a leading platform for *in vivo* gene therapy, highlighted by recent clinical successes such as Luxturna (for inherited retinal dystrophy), Zolgensma (for spinal muscular atrophy), and Roctavian and Hemgenix (for hemophilias A and B, respectively) (Kohn et al., 2023; Park et al., 2026). While the viral capsid provides some degree of cell-type specificity through tropism, the packaged promoter is the critical determinant of the spatial distribution and expression level of the transgene (D. Wang et al., 2019). Optimizing promoters for precise control of transgene expression is therefore essential to maximize gene therapy safety and efficacy. The ideal promoter for rAAV applications must satisfy two main criteria. First, due to the limited DNA payload capacity of rAAV vectors (approximately 4.7 kilobases), it must be compact to accommodate the transgene (C. Li & Samulski, 2020). Second, it must drive cell-type specific (*i.e*., on-target) expression at endogenous-like levels to mitigate adverse reactions associated with ectopic expression or dose-dependent transgene toxicity (Alstyne et al., 2021; Xiong et al., 2019). Beyond these two requirements, it is also highly desirable if the promoter exhibits cross-species functional conservation to ensure that validation in preclinical models, such as mice and non-human primates, translates reliably to human patients (Galvan et al., 2026; Gomes et al., 2024; Jüttner et al., 2019).

Despite the need for optimized rAAV promoters, designing sequences that simultaneously satisfy all three criteria remains challenging. Traditional rational design strategies include utilizing the proximal regions of endogenous on-target promoters (Bainbridge et al., 2008), concatenating relevant transcription factor (TF) motifs (Hwang et al., 2005), and assembling conserved *cis*-regulatory elements (CREs) from genes with desirable expression profiles (Fornes et al., 2023). However, these approaches often require compromising one criterion for another and cannot universally produce satisfactory solutions for all cell types. Sequence-to-function deep learning models have emerged as powerful tools for decoding the complex *cis*-regulatory logic of the genome (Avsec et al., 2021, 2026; Linder et al., 2025) (reviewed in (Eraslan et al., 2019)). When trained on cell-type-specific chromatin accessibility, these models learn to predict the probability that a given sequence is accessible within that cell type, and can then be computationally interrogated to identify the underlying features, such as relevant TF motifs, that led to the prediction (Novakovsky et al., 2023). Notably, models trained on accessibility data can also serve as “oracles” to guide the design of CREs with desired expression characteristics (Almeida et al., 2024; Taskiran et al., 2024; S. K. Wang et al., 2026). For example, they can be coupled with *in silico* directed evolution (Lal et al., 2025) to iteratively mutate and select sequences that maximize the gap between on-target and off-target predictions (*i*.*e*., MinGap) (Gosai et al., 2024). Alternatively, gradient-based editing approaches, such as Ledidi (Schreiber et al., 2025), can directly optimize DNA sequences by backpropagating desired properties through the model. Because both strategies rely on satisfying the learned *cis*-regulatory grammar (rather than sequence homology), they generate novel sequences unconstrained by evolutionary divergence, enabling the creation of highly compact CREs. However, there is no guarantee that CREs designed in this manner to maximize specificity in one species will exhibit cross-species functional conservation when tested *in vivo*.

Here, we introduce a generalizable deep learning framework for the design of CREs with high on-target specificity and cross-species functional fidelity. We evaluated the framework by targeting Müller glia (MG), a retinal glial population that provides critical structural and metabolic support to neurons (Goldman, 2014). In lower vertebrates such as zebrafish, MG can regenerate all retinal neuronal subtypes upon injury, including photoreceptors. However, mammalian MG lack this innate capacity and require additional reprogramming steps to drive neuronal commitment, such as the forced expression of the proneural TF ASCL1 (Hoang et al., 2020; Martin & Poché, 2019). This latent neurogenic potential makes MG compelling targets for gene therapy, particularly for regenerative strategies to replace lost photoreceptors in blinding diseases such as age-related macular degeneration (Goldman, 2014).

To establish a robust foundation for model training and cross-species validation, we first compiled a comprehensive single-cell chromatin accessibility atlas of the human retina (Orozco et al., 2023) and generated a corresponding one for the mouse retina. We then fine-tuned deep learning models on these atlases to separately predict retinal cell-type-specific accessibility in both species. Using two complementary strategies, we designed compact, highly specific CREs targeting MG and mechanistically mapped their specificity to the creation of motifs for the TF LHX2. Finally, we demonstrated that the designed CREs drive highly restricted MG-specific expression in the murine retina *in vivo*, highlighting the power of deep learning to engineer precision gene therapy tools.

## Results

### Cell-type resolution accessibility atlases of the mammalian retina

To establish a robust foundation for training our deep learning models, we compiled comprehensive single-cell (sc) and single-nucleus (sn) chromatin accessibility atlases of the mammalian retina. Specifically, we leveraged snATAC-seq data from the human retina, retinal pigment epithelium (RPE), and choroid (7 donors; 82,186 cells), comprising 317,515 consensus chromatin accessibility peaks across 17 cell types (Orozco et al., 2023). Because Müller glia (MG) exhibited high co-accessibility with astrocytes (Pearson *R* = 0.86, Bonferroni-corrected *p* < 0.001; **Fig. S1A**), we complemented this data with human brain astrocyte sc/snATAC-seq data from two additional studies to better model these distinct but closely related glial populations (Corces et al., 2020; Morabito et al., 2021).

Validating human *cis*-regulatory elements (CREs) in model organisms remains challenging (Jüttner et al., 2019). To overcome this limitation and enable cross-species comparison, we established a reference atlas of chromatin accessibility in the mouse retina. We performed sn-multiome sequencing, jointly profiling gene expression (RNA-seq) and chromatin accessibility (ATAC-seq) in 54,073 cells from C57BL/6J mice (**Methods**). Cell-type identities were annotated using label transfer from the Mouse Retinal Cell Atlas (J. Li et al., 2024). Cell types formed distinct, well-defined clusters in both the RNA (t-SNE) and ATAC (UMAP) embeddings (**Fig. 1A**), confirming that cell-type identity was preserved across modalities and that the multiome data captured the known cellular diversity of the mouse retina. To assess the correspondence between these modalities, we further examined the expression and chromatin accessibility of well-established retinal marker genes (**Fig. 1B**). Cell-type-specific markers exhibited high concordance between RNA expression and gene activity scores. For instance, the MG marker *Rlbp1* was enriched in both RNA expression and chromatin accessibility exclusively within the MG population.

**Figure 1:**
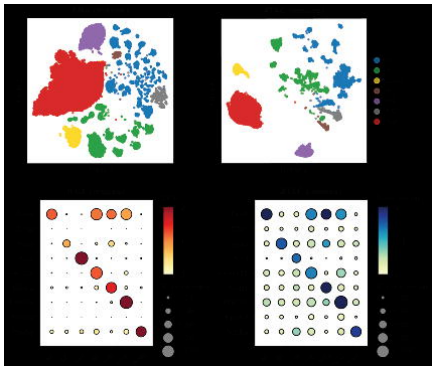
Overview of the mouse retinal multiome data. (**A**) Visualization of the mouse retinal single-nucleus multiome data using TSNE for RNA expression (left) and UMAP for ATAC chromatin accessibility (right). Cells are colored by their assigned cell type from the Mouse Retinal Cell Atlas (J. Li et al., 2024). (**B**) Dot plots comparing gene expression (RNA, left) and ArchR-derived gene activity scores (Granja et al., 2021) (ATAC, right), which provide a gene-centric measure of chromatin accessibility, for canonical marker genes across identified cell types. AC, amacrine cells; BC, bipolar cells; HC, horizontal cells; MG, Müller glia; RGC, retinal ganglion cells.

Across the mouse dataset, we identified a total of 464,996 consensus chromatin accessibility peaks. Consistent with the human dataset, the majority of peaks were distal (90.1%) rather than located in promoter regions (<1 kb from a transcription start site; 9.9%). Consistent with the human dataset, the mouse dataset was dominated by rods (n = 20,157), followed by amacrine cells (AC; n = 9,395) and bipolar cells (BC; n = 8,419), with MG comprising 7.2% of the total cells (n = 3,909). To better compare the retinal accessibility landscapes across species, we performed a synteny analysis between the human and mouse peak sets (**Methods**). A considerable fraction of the human (22.9%) and mouse (33.8%) peaks had an orthologous peak in the other species, although this varied across individual cell types. For example, only 12.8% of peaks identified in human horizontal cells (HC) had an orthologous peak in mouse HCs, compared to 58.6% when mapping in the opposite direction, an asymmetry that likely reflects the low number of these cells captured in the mouse dataset (n = 376). Differentially accessible (DA) peaks showed even lower overall conservation, averaging only 8.8% cross-species-mappability across human cell types compared to 12.3% across mouse cell types (**Fig. S1B**).

### Deep learning design and in silico validation of retinal CREs

We developed a three-step deep learning workflow for the design and *in silico* validation of cell-type-specific retinal CREs, consisting of model training, sequence optimization, and cross-model validation (**Fig. 2A**).

**Figure 2:**
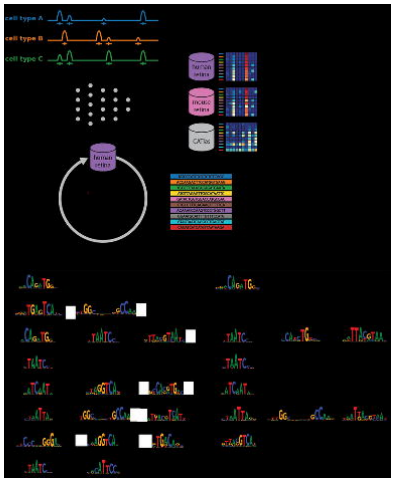
Deep learning framework for the design and in silico validation of cell-type-specific retinal cis-regulatory elements. (**A**) Schematic of the deep learning workflow. Binarized pseudobulk peak matrices are used to fine-tune a custom Enformer architecture initialized with the original model weights (Avsec et al., 2021). Starting sequences are iteratively optimized using the human model coupled with directed evolution (Lal et al., 2025) or Ledidi (Schreiber et al., 2025) to maximize the gap between their predicted on-target and off-target accessibility in the human retina (*i*.*e*. MinGap; (Gosai et al., 2024)). Evolved elements undergo *in silico* validation across the human and mouse retinal models, as well as a model trained on the CATlas dataset (Zhang et al., 2021), to prioritize candidates with high predicted on-target retinal accessibility, cross-species functional conservation, and minimal off-target accessibility in non-retinal cell types and tissues. (**B**) Transcription factor (TF) motifs discovered by TF-MoDISco (Shrikumar et al., 2018) in the human and mouse models for each retinal cell type. Discovered motifs are annotated by comparison to non-redundant TF motif clusters derived from the JASPAR 2024 database (Rauluseviciute et al., 2023) using TOMTOM (Schreiber, 2025b). AC, amacrine cells; BC, bipolar cells; HC, horizontal cells; MG, Müller glia; RGC, retinal ganglion cells; RPE, retinal pigment epithelium. The asterisk (*) denotes the three human astrocyte peak sets (one retinal and two brain-derived). Photoreceptors are denoted as “Cone+Rod”.

#### Step 1: Model training

Prior to training, we pseudobulked the single-cell retinal atlases at a 500-base-pair (bp) resolution per cell type for human (317,515 peaks × 19 cell types, including three different astrocyte tracks: one retinal and two brain-derived) and mouse (464,996 peaks × 7 cell types), resulting in a binarized matrix denoting peak accessibility status (1 for “open” and 0 for “closed”) for each species. We then fine-tuned a custom Enformer-based architecture on each pseudobulk matrix, initializing it with weights from the original model but reducing it to a single transformer layer (Avsec et al., 2021), to predict, for each cell type, the probability that a sequence represents an accessible peak (**Methods**). The resulting species-specific models exhibited robust performance across all cell types (**Fig. S2**). To verify that they had captured comparable retinal *cis*-regulatory grammars rather than species-specific sequence biases or technical artifacts, for each cell type, we applied TF-MoDISco (Shrikumar et al., 2018) to identify high-importance transcription factor (TF) motifs based on model-derived attribution scores (**Fig. 2B**). Discovered motifs were subsequently annotated by comparing them against a non-redundant set of TF motifs (**Methods**). Notably, despite the previously observed low genomic synteny of DA peaks (averaging only 9% to 12% cross-species conservation), the human and mouse models independently learned nearly identical *cis*-regulatory grammars, with the same TF motifs enriched in corresponding cell types across both species (**Table S1**). For instance, CRX-like motifs in photoreceptors (Chen et al., 1997; Furukawa et al., 1997), ONECUT in HCs (Klimova et al., 2015; Wu et al., 2013), and LHX in MG (Melo et al., 2016) were consistently identified as top predictive features in both models.

#### Step 2: Sequence optimization

For each cell type of interest, including MG, 10 DA peaks were randomly selected as starting sequences and iteratively optimized using the human model to maximize their predicted accessibility in that cell type (*i*.*e*., on-target) while minimizing predicted accessibility across all other cell types (*i*.*e*., off-target). We used two complementary design strategies: directed evolution (Lal et al., 2025), which introduces point mutations and selects for improved predicted cell-type-specific accessibility over successive iterations, and Ledidi (Schreiber et al., 2025), a gradient-guided sequence editing approach. Both strategies used the human retinal model and shared the same objective function: maximizing the difference between predicted on-target and off-target accessibilities (*i*.*e*., MinGap (Gosai et al., 2024)).

#### Step 3: Cross-model validation

The designed CREs underwent *in silico* validation across three independent models: the human and mouse retinal models, and a multi-tissue chromatin accessibility model spanning 204 different human cell types and tissues from CATlas (Zhang et al., 2021) (**Methods**). Specifically, they were assessed for their predicted on-target accessibility in both retinal models, as well as their predicted non-retinal accessibility in the CATlas model.

We also developed a complementary approach using a human regression model trained on pseudobulked fragment counts scaled between 0 and 1 to predict normalized ATAC signal rather than binary peak accessibility status (**Note S1**).

### Designed CREs exhibit high predicted MG-specific accessibility and minimal off-target accessibility

We applied both directed evolution and Ledidi to design MG-specific CREs. As expected, starting sequences derived from endogenous DA MG peaks already exhibited substantial on-target accessibility in both the human and mouse models (**Fig. 3A**). Using directed evolution, sequences reached near-maximum predicted MG-specific accessibility within the first 10 iterations (MinGap score = 0.94, where 1 represents the maximum), after which per-iteration gains plateaued (**Fig. 3B**). This rapid convergence suggests that a relatively small number of point mutations is sufficient to create cell-type-specific CREs, consistent with recent findings (Taskiran et al., 2024). CREs designed using Ledidi achieved comparable MG-specific accessibility (**Fig. 3A**). Moreover, evaluation of the best-evolved designs (*i*.*e*., highest-MinGap-scoring CREs evolved from each starting sequence) across the 204 cell types and tissues in the CATlas model confirmed that both strategies produced CREs with minimal predicted off-target accessibility (**Fig. 3C**), indicating that they are highly specific for MG rather than broadly accessible. Similar rapid evolutionary trajectories and high on-target accessibility levels were observed when evolving cell-type-specific CREs for other retinal cell types or when starting the design process from random sequences (**Figs. S3** and **S4**). However, the consistently lower final cell-type specificity of the CREs designed from random sequences, as measured by MinGap, suggests that partially on-target endogenous sequences provide a better starting point for maximizing cell-type specificity.

**Figure 3:**
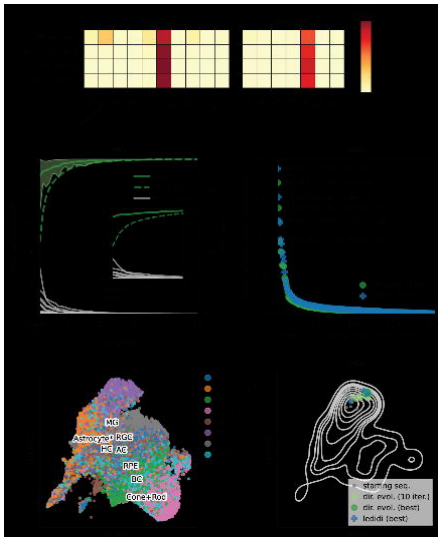
In silico validation of Müller glia (MG)-specific cis-regulatory elements. (**A**) Heatmaps showing the average predicted chromatin accessibility in the human and mouse retinal models for the starting sequences (starting seqs.) and best-evolved elements (*i*.*e*., the highest-MinGap-scoring elements evolved from each starting sequence, where MinGap is defined as the difference between the minimum predicted on-target and maximum predicted off-target accessibility in the human retina (Gosai et al., 2024)) using directed evolution (dir. evol.) or Ledidi. For directed evolution, partially-evolved elements at iteration 10 (10 iter.) are also shown. (**B**) Average predicted chromatin accessibility (shaded area indicates the min-max range) and MinGap score (secondary y-axis) in the human retinal model across the 50 iterations of directed evolution. The inset highlights the rapid gain in predicted MG accessibility and specificity (*i*.*e*., MinGap) within the first 10 iterations. (**C**) Predicted chromatin accessibility for the best-evolved elements across 204 cell types and tissues from the CATlas model (*i*.*e*., off-target), ranked by accessibility in descending order. (**D**) UMAP visualization of approximately 2,500 differentially accessible (DA) peaks from selected major cell classes in the multiome data, embedded using the human model’s latent space. (**E**) Projection of starting MG sequences and evolved elements onto the MG DA peak landscape. AC, amacrine cells; BC, bipolar cells; HC, horizontal cells; MG, Müller glia; RGC, retinal ganglion cells; RPE, retinal pigment epithelium. One asterisk (*) denotes the three human astrocyte peak sets (one retinal and two brain-derived); two asterisks (**) denote non-retinal peak sets (*e*.*g*., B and T cells). Photoreceptors are denoted as “Cone+Rod”.

To visualize the relationship between designed CREs and endogenous sequences, we embedded approximately 2,500 DA peaks from ACs, astrocytes, BCs, HCs, MG, photoreceptors, retinal ganglion cells (RGC), and RPE into the human retinal model’s latent space (**Fig. 3D**). The resulting UMAP preserved distinct cell type territories, with MG and astrocyte peaks clustering in close proximity, reflecting their *cis*-regulatory similarities. We then projected the starting MG sequences, partially-evolved CREs (10 iterations of directed evolution), and best-evolved designs onto this landscape, confirming that both directed evolution and Ledidi progressively push CREs toward the most extreme region of the MG-specific territory (**Fig. 3E**). This optimization trajectory suggests that the designed CREs are biologically plausible, occupying a sequence space consistent with known MG biology (Lal et al., 2024).

### Directed evolution and Ledidi achieve MG-specific accessibility by creating LHX2 motifs

To understand the mechanistic basis for the designed MG-specific accessibility, we analyzed the enrichment of TF motifs in the designed CREs relative to the endogenous set of 2,500 DA MG peaks (**Fig. 4A**; **Methods**). The LHX motif (C100:HOXB_LHX) emerged as the most enriched during the design process, while the RREB1 motif (C131:RREB1) was the most depleted. This pattern was consistent across both design strategies, and was already apparent after only 10 iterations of directed evolution, suggesting that the LHX motif is the primary determinant of MG-specific accessibility. Similar enrichments of relevant cell-type-specific TF motifs were observed when designing DA CREs for other retinal cell types (**Fig. S5**), such as ONECUT in HCs and TEAD in RPE, the latter of which serves as a critical regulator of RPE identity and homeostasis in conjunction with co-activators YAP and TAZ (Miesfeld et al., 2015).

**Figure 4:**
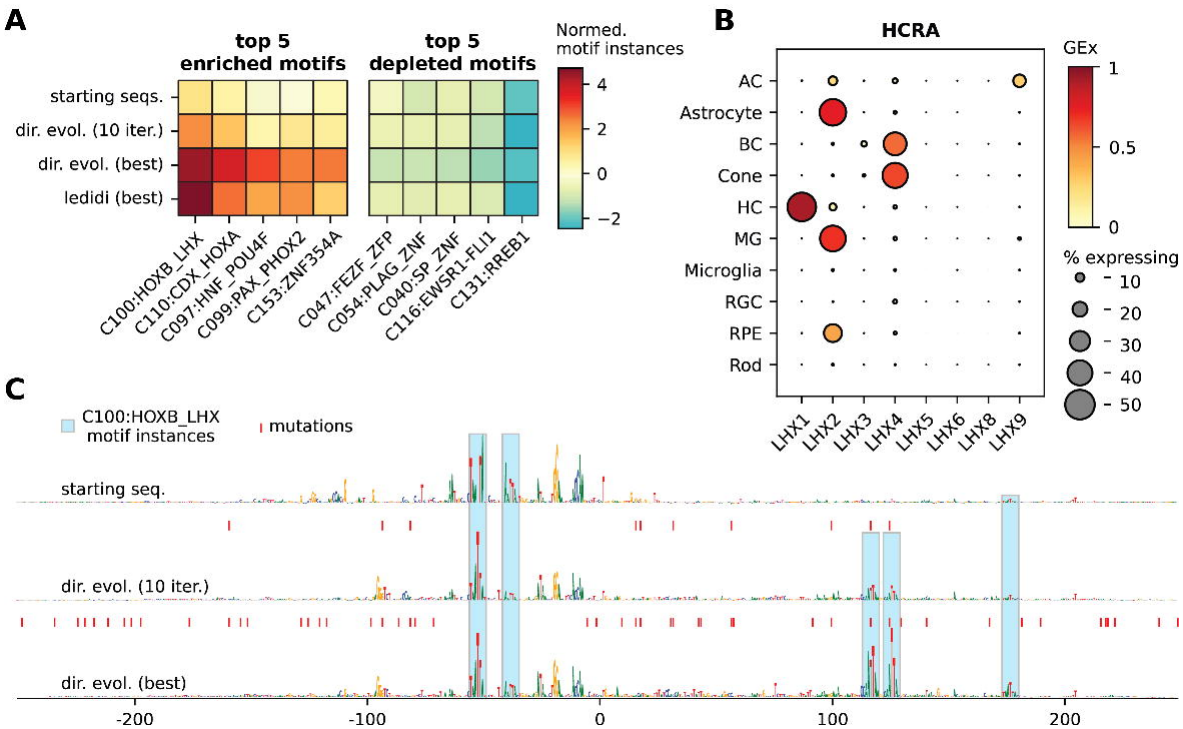
Enrichment of the LHX2 motif in Müller glia (MG)-specific cis-regulatory elements. (**A**) Heatmaps showing the average normalized counts of the top 5 enriched (left) and depleted (right) transcription factor (TF) motifs for the starting Müller glia (MG) sequences (starting seqs.) and best-evolved elements (*i*.*e*., the highest-MinGap-scoring elements evolved from each starting sequence, where MinGap is defined as the difference between the minimum predicted on-target and maximum predicted off-target accessibility in the human retina (Gosai et al., 2024)) using directed evolution (dir. evol.) or Ledidi relative to approximately 2,500 differentially accessible MG peaks. For directed evolution, partially-evolved elements at iteration 10 (10 iter.) are also shown. (**B**) Dot plot comparing the expression of *LHX* family genes in the Human Retinal Cell Atlas (HRCA). (**C**) Sequence optimization of a representative MG element using directed evolution. Attribution scores, representing the importance of each nucleotide for predicted MG accessibility, are shown as sequence logo heights. Red ticks along the x-axis indicate point mutations introduced relative to the starting sequence, and blue shaded regions denote identified C100:HOXB_LHX motif instances. AC, amacrine cells; BC, bipolar cells; HC, horizontal cells; MG, Müller glia; RGC, retinal ganglion cells; RPE, retinal pigment epithelium.

Indeed, examination of gene expression data from the Human Retinal Cell Atlas (J. Li et al., 2026) confirmed that *LHX2* is the only LHX family member expressed at high levels in MG, with elevated expression also observed in astrocytes and marginal expression in the RPE (**Fig. 4B**). LHX2 is a TF with well-established roles in retinal development and MG biology. In the murine retina, *Lhx2* is required for MG development, regulating the proliferation of gliocompetent retinal progenitors, activating MG-specific gene expression, and driving the terminal differentiation of MG morphological features (Melo et al., 2016). Interestingly, recent evidence suggests that LHX2 also suppresses astrocyte proliferation in the cerebral cortex (Iyer et al., 2025), supporting its context-dependent role in our models: driving MG accessibility while remaining relatively unimportant in astrocytes (**Fig. 2B**). Notably, *in silico* experiments supported both the requirement and sufficiency of the LHX2 motif for driving MG-specific accessibility (**Methods**). When we constrained the design process by penalizing the creation of LHX2 motifs into random sequences, we failed to evolve CREs toward MG accessibility (**Fig. S6A**). Conversely, inserting LHX2 motifs (instead of point mutations) was sufficient to design CREs with predicted MG-specific accessibility comparable to those designed via directed evolution or Ledidi starting from endogenous sequences (average MinGap score = 0.99; **Fig. S6B**). However, they occupied a distinct region of the human retinal model’s latent space that neither overlapped with DA MG peaks nor CREs designed from endogenous sequences (**Fig. S6C**), suggesting that LHX2-motif insertion alone results in non-natural designs.

Next, we performed sequence-level attribution analysis of selected designed CREs (**Figs. 4C** and **S7**). Attribution scores, which quantify the contribution of each nucleotide to the model’s predicted output, revealed that the newly created LHX2 motifs became highly important. Moreover, the design process actively avoided disrupting any existing LHX2 sites in the starting sequences.

### Designed CREs drive highly specific MG expression in vivo

We evaluated a panel of candidate CREs (**Table S2**; **Methods**) for their ability to drive MG-specific GFP expression in the murine retina using rAAV-mediated reporter assays, including DA MG peaks lacking predicted LHX motifs, partially- and fully-evolved CREs, and CREs designed using a human regression model (**Note S1**). Quantification of reporter expression revealed that one of the partially-evolved CREs (MG #10) achieved higher MG specificity than a minimal version of the hRGR promoter (**Figs. 5A-D**), which served as a benchmark given its comparable compactness (709 bp) and restricted MG expression in the murine retina when delivered via rAAV (Jüttner et al., 2019). Notably, one of the regression model-designed CREs (MG #6) also exhibited higher MG specificity than the hRGR promoter. Specifically, the mean percentage of GFP+ cells co-expressing the MG marker Sox9 was 40-60% for these CREs, compared to approximately 20% for hRGR (**Fig. 5E**). In addition, the hRGR promoter exhibited substantial off-target GFP+ cell labeling in the ganglion cell layer, an observation not reported in Jüttner *et al*., likely because they evaluated a longer version (2,000 bp) of the mouse promoter rather than the human minimal version that we used.

**Figure 5:**
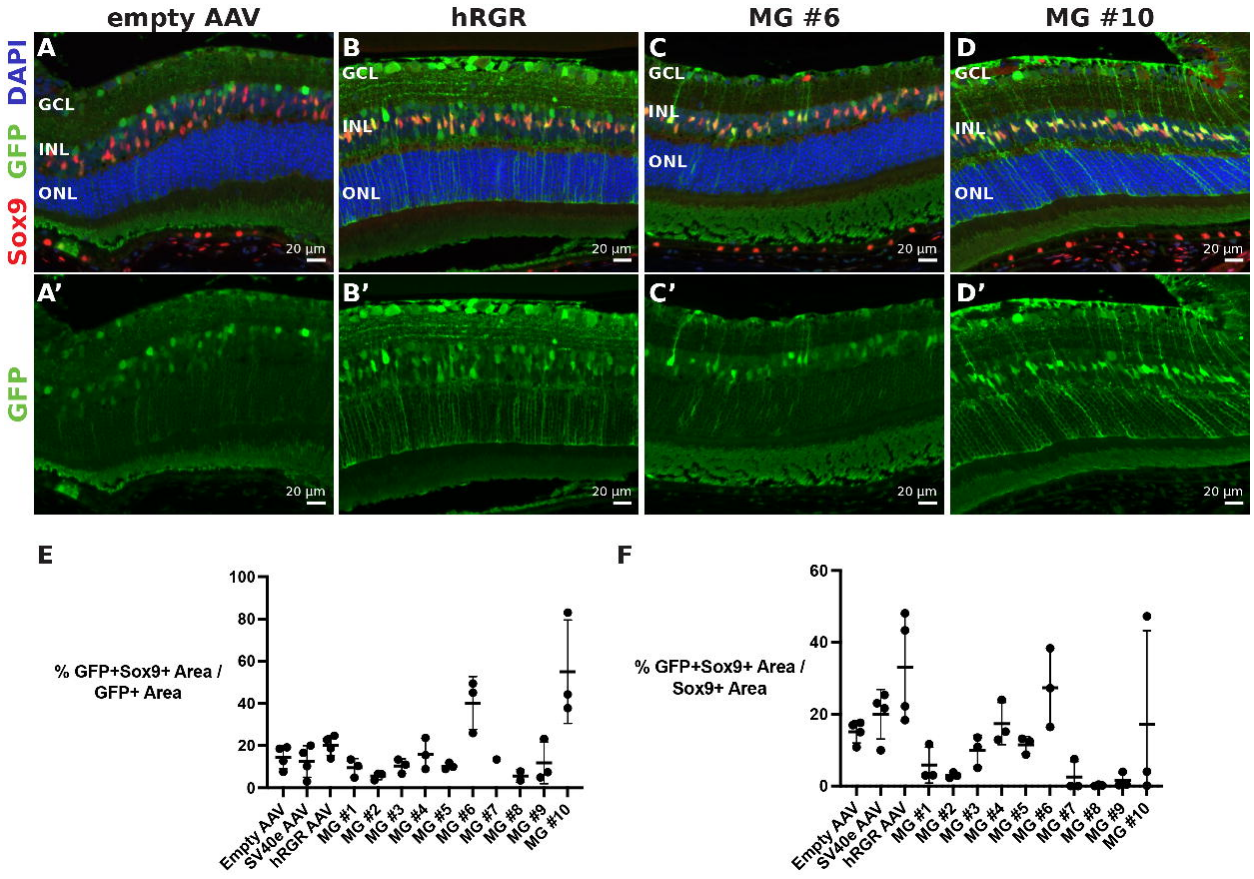
Experimental validation of Müller glia (MG)-specific elements in the murine retina. (**A-D**) Confocal images of adult mouse retinal sections 5 weeks post-intravitreal injection with rAAV vectors carrying GFP under the control of an empty vector (minP only; **A**), the hRGR promoter (**B**), or designed MG elements #6 (**C**) and #10 (**D**). Sections are stained for GFP (green), the MG marker Sox9 (red), and DAPI (blue). (**A’-D’**) Isolated GFP channel highlighting the morphology and distribution of transduced cells. (**E**) Quantification of specificity, defined as the percentage of GFP+ area that is Sox9+. (**F**) Quantification of transduction efficiency, defined as the percentage of the total Sox9+ area that is GFP+. Data points denote individual biological replicates; error bars indicate the mean ± standard deviation. DAPI, 4’,6-diamidino-2-phenylindole; GFP, green fluorescent protein; GCL, ganglion cell layer; INL, inner nuclear layer; ONL, outer nuclear layer.

The mean transduction efficiency of MG #6 and MG #10 was comparable to the hRGR promoter (**Fig. 5F**), with approximately 20-30% of the total Sox9+ MG population labelled compared to approximately 35% for hRGR. However, group sizes were small, and transduction efficiency varied substantially within groups. For example, only the eyes of one of three animals injected with MG #10 exhibited robust GFP labeling *in vivo*. We attribute this variability to technical factors, such as differences in injection quality and section sampling, rather than intrinsic differences among the CREs tested. While larger cohorts would provide more definitive estimates of transduction efficiency, these results demonstrate that the designed CREs are more on-target, achieving superior MG specificity without substantially compromising penetrance, validating the capacity of our deep learning-driven sequence optimization framework for engineering precise *in vivo* gene delivery tools.

## Discussion

The development of safe and effective rAAV-based gene therapies has historically been bottlenecked by the need for optimized promoters that are both compact and cell-type specific (*i*.*e*., on-target). Moreover, cross-species functional conservation, a critical feature for clinical translation, has remained largely underexplored in promoter engineering. Traditional rational design often forces a compromise between payload capacity and on-target specificity. Here, we demonstrate that sequence-to-function deep learning models can circumvent these limitations by guiding the design of *cis*-regulatory elements (CREs) that satisfy all three criteria. By separately modeling accessibility in the human and mouse retinas, we designed CREs targeting Müller glia (MG) that are both compact and predicted to maintain on-target specificity across the two species (**Fig. 3A**).

Because designed CREs are not constrained by evolution, they can sample a broader combinatorial sequence space to optimize *cis*-regulatory logic. As evidenced in the retinal embeddings (**Fig. 3E**), we designed extreme versions of differentially accessible (DA) MG peaks through the creation of novel instances of the LHX2 transcription factor (TF) motif. While LHX2 is a well-established regulator of MG (Melo et al., 2016), it has also been proposed to act as a transcriptional repressor in astrocytes (Iyer et al., 2025), a cell type that exhibited high co-accessibility with MG (**Fig. S1**). By creating these motifs, the sequence optimization process converged on a multi-faceted strategy: actively increasing on-target accessibility in MG while potentially repressing a closely related glial population (*i*.*e*., astrocytes) and penalizing off-target accessibility in other retinal cell types. This specificity extended beyond the local retinal environment: when evaluated against an independent whole-body chromatin accessibility model trained on the CATlas dataset (Zhang et al., 2021), the designed CREs exhibited minimal predicted accessibility across hundreds of non-retinal human cell types and tissues. Notably, *in vivo* validation demonstrated that the designed CREs significantly outperformed the endogenous hRGR benchmark in the murine retina, achieving highly restricted MG-specific expression without substantially compromising transduction penetrance.

Mechanistic interpretability proved critical for validating the designed CREs prior to experimental validation. By interrogating them through our models, we confirmed that both directed evolution and Ledidi achieved high predicted on-target accessibility by convergently evolving motifs for known biologically relevant TFs (*e*.*g*., LHX2 in MG or ONECUT in horizontal cells). We also identified practical constraints for designing CREs. While the design process can theoretically start from random DNA, doing so forces predictions into uncharacterized sequence spaces that were not represented in the model’s training data (**Fig. S6C**). Instead, starting from endogenous DNA that already partially exhibits the desired specificity anchors the design process within the model’s known biological context, producing CREs that are both more specific (**Figs. S3** and **S4**) and more biologically realistic (Lal et al., 2024). When starting from endogenous sequences, just 10 iterations were sufficient for achieving high predicted on-target accessibility, serving as a practical threshold that minimizes deviation from natural biological sequences.

Establishing a minimal mutagenesis threshold is important to mitigate the out-of-distribution risks associated with generative design (Lal et al., 2024). Extensive sequence manipulation, such as evolving sequences for up to 50 iterations or introducing excessive mutations via Ledidi (Schreiber et al., 2025), delves into unexplored territory outside the original training data and can result in confident but biologically unviable computational artifacts that fail when tested *in vivo*. We observed this when designing CREs using regression models trained on scaled fragment counts (**Note S1**). For instance, when applying MinGap to maximize the difference between predicted on-target and off-target accessibility (Gosai et al., 2024), the design process artificially inflated accessibility across all cell types: on-target predictions increased to levels two to three times higher than the expected biological maximum, while off-target accessibility levels also increased. Alternatively, applying a ratio-based objective function, where on-target accessibility was divided by the maximum predicted off-target accessibility, simply minimized the off-target denominator and failed to optimize for on-target accessibility. In contrast, optimizing against binary accessibility (open or closed) proved far more effective. DA peaks typically display intermediate accessibility levels, whereas peaks with maximum accessibility tend to be broadly accessible across cell types. By binarizing accessibility, we forced the design process to prioritize the probability of being accessible on-target rather than inflating ubiquitous accessibility, ensuring the designed CREs remained highly specific and biologically plausible.

To further improve cross-species functional conservation, future iterations of this work could leverage a multi-head model architecture to simultaneously learn human and mouse retinal accessibility. By using a shared encoder with species-specific decoder heads, such a model could capture unified representations of *cis*-regulatory logic while avoiding species-specific features such as repeats. Alternatively, we could apply a joint optimization function that simultaneously applies MinGap to the outputs from both the human and mouse models. However, as we observed for regression-based designs, such joint approaches require careful calibration to avoid artificial inflation of accessibility values. Ultimately, a balanced multi-objective framework that preserves the advantages of binarization while jointly constraining on-target probability and specificity across species could enhance evolutionary convergence toward genuinely conserved CREs rather than species-specific solutions.

A current limitation of our framework, and of similar approaches, is the use of chromatin accessibility as a proxy for gene expression (Almeida et al., 2024; Taskiran et al., 2024; S. K. Wang et al., 2026). Biologically, only a fraction of accessible regions predicted to act as enhancers successfully drive robust expression in massively parallel reporter assays (MPRAs) (Kwasnieski et al., 2014). However, CREs designed using sequence-to-function models drive significantly stronger and more specific expression levels than the endogenous accessible sequences from which they originate (Gosai et al., 2024). While future models will benefit from training directly on multi-tissue MPRA data, our results confirm that models trained on comprehensive single-cell accessibility atlases are already capable of designing highly on-target CREs.

An additional technical limitation of our study was intrinsic to the dual-cassette rAAV construct used for the mouse *in vivo* validation. To evaluate the designed CREs, they were cloned upstream of a minimal promoter (minP) containing a TATA-box motif to initiate transcription driving eGFP. A separate constitutive CAG promoter was included on the same vector to drive *Cre* recombinase. In practice, the baseline minP control exhibited leaky eGFP expression and the CAG promoter negligible Cre expression, which could indicate promoter or transcriptional interference, a well-documented phenomenon in multi-promoter rAAV vectors where adjacent CREs can cause promoter hijacking, steric hindrance, or transcriptional suppression (Foti et al., 2009). While the restricted spatial specificity driven by our designed CREs remained clearly discernible above this background, these vector-level artifacts underscore the importance of using streamlined, single-promoter architectures or incorporating insulating elements in future *in vivo* reporter assays.

Taken together, the proposed framework demonstrates the utility of generative deep learning for engineering precision gene therapy tools. By bypassing the limitations of evolutionary divergence and traditional rational design, it enables the rapid generation of highly compact, cell-type-specific, and cross-species functionally conserved CREs.

## Methods

### Single-cell multiome sequencing

Eight retinas from 11- and 93-week-old C57BL/6 mice (4 per age group) were rapidly dissected from enucleated eyes in cold PBS, immediately snap-frozen on dry ice, and stored at -80°C until further processing. Snap-frozen retinal tissue was transferred into a glass Dounce homogenizer containing 500 ⊠L chilled 0.1× lysis buffer (10 mM Tris-HCl, pH 7.4, 10 mM NaCl, 3 mM MgCl⊠, 0.01% Tween-20, 0.01% Nonidet P40 Substitute, 0.001% digitonin, 1% BSA, 1 mM DTT, 1× RNase inhibitor in nuclease-free water) and immediately homogenized. Homogenates were incubated on ice for 5 minutes, gently triturated using a wide-bore pipette tip, and incubated on ice for an additional 10 minutes. Following lysis, 500 ⊠L of chilled wash buffer (10 mM Tris-HCl, pH 7.4, 10 mM NaCl, 3 mM MgCl⊠, 1% BSA, 1 mM DTT, 1× RNase inhibitor in nuclease-free water) was added, and the samples were mixed gently by pipetting. Nuclei suspensions were sequentially filtered through 70 ⊠m and 40 ⊠m Flowmi cell strainers to remove debris and aggregates. Filtered nuclei were centrifuged at 500 × *g* for 5 minutes at 4°C, washed twice with chilled wash buffer, and resuspended by gentle pipetting. Nuclei concentration was determined using a Countess II FL Automated Cell Counter. Nuclei were subsequently pelleted and resuspended in chilled diluted nuclei buffer (1× Nuclei Buffer with 1 mM DTT and 1× RNase inhibitor in nuclease-free water). Samples were maintained on ice throughout the procedure and processed immediately for downstream single-nucleus multiome library preparation.

### Multiome library preparation and sequencing

Single-nucleus (sn) ATAC and gene expression libraries were constructed using the Chromium Next GEM Single Cell Multiome ATAC + Gene Expression kit (10x Genomics) according to the manufacturer’s protocols. Ten thousand nuclei were targeted for library preparation. Each nuclei suspension was submitted for library preparation using the Chromium Next GEM Chip J Single Cell Kit (10X Genomics), following the manufacturer’s instructions. The resulting libraries were sequenced on an Illumina NovaSeq S4 targeting 25,000 read pairs per nucleus.

### Multiome read alignment

Sequencing reads from the sn-multiome libraries were mapped to the mouse reference genome (GRCm38) using Cell Ranger ARC (version 2.0.0). The resulting raw gene expression matrices were imported into R (version 4.5.2) and converted to the SingleCellExperiment format using the Bioconductor ecosystem (version 3.22) (Huber et al., 2015).

### snRNA-seq data processing and analysis

Downstream processing and analysis of the sn-multiome RNA component were performed using scrapper (version 1.4.0) (Lun & Kancherla, 2023). Specifically, we computed per-cell quality control (QC) metrics, including total UMI count, the number of detected genes, and mitochondrial read proportions (computeRnaQcMetrics). Low-quality cells were filtered out using an outlier-based strategy of 3 median absolute deviations from the median, applied independently to each metric (suggestRnaQcThresholds). Counts were then normalized using library-size scaling factors and subsequently log-transformed (normalizeCounts). Highly variable genes (HVGs) were identified by fitting a mean-dependent trend to the per-gene log-expression variances, retaining the top 2,000 genes with the largest residuals (modelGeneVariances). Dimensionality reduction was subsequently performed via principal component analysis (PCA) on the HVG log-expression matrix, retaining the top 25 principal components (runPca). Finally, shared nearest-neighbor graph construction (k = 10), Leiden community detection (multilevel method), t-SNE (perplexity = 30), and UMAP (15 neighbors, min-dist = 0.1) embeddings were jointly computed from the PCA space (runAllNeighborSteps).

### Cell-type annotation using label transfer

Cell-type identities were transferred from the Mouse Retinal Cell Atlas (MRCA) (J. Li et al., 2024) using Seurat’s anchor-based approach (version 5.4.0) (Hao et al., 2024). Cluster identities were subsequently assigned manually by inspecting MRCA prediction scores and cell-type proportions per cluster, resulting in the following groups: amacrine cells (AC), bipolar cells (BC), cones, horizontal cells (HC), Müller glia (MG), retinal ganglion cells (RGC), and rods. Clusters that could not be confidently assigned were regarded as putative doublets for downstream removal.

### snATAC-seq data processing and analysis

Downstream processing and analysis of the sn-multiome ATAC component were performed using ArchR (version 1.0.3.1) (Granja et al., 2021). Specifically, Arrow files were generated from the aligned fragments and used to initialize an ArchR project. Doublets were identified and removed using filterDoublets, and any cells lacking matched RNA data were excluded. Dimensionality reduction was performed separately on the ATAC tile matrix and the gene expression matrix via iterative latent semantic indexing (LSI; addIterativeLSI), and the resulting embeddings were concatenated (addCombinedDims). Cell clusters were then identified within the combined LSI space using a graph-based approach (addClusters), and a UMAP embedding was computed for visualization (addUMAP). Pseudobulk replicates were generated for each cluster (addGroupCoverages), with the analysis restricted to autosomes and the X chromosome. Peak calling was subsequently performed using addReproduciblePeakSet with MACS2 (version 2.2.9.1) (Feng et al., 2012), setting the effective genome size to 1.87 × 10⊠ base pairs (bp) (for the mm10 reference), resulting in a peak-by-cell count matrix (addPeakMatrix). Finally, cell-type identities were transferred from the snRNA-seq analysis by intersecting barcodes present in both modalities. ATAC clusters were annotated manually, informed by these RNA-derived cell-type assignments, resulting in the same cell types (AC, BC, cones, HC, MG, RGC, and rods). Cells within clusters that could not be confidently annotated were regarded as putative doublets and removed.

### Generation of pseudobulk peak matrices

We generated cell-type-resolution pseudobulk peak matrices as follows. For mouse, per-cell-type pseudobulk fragment files were extracted from the ArchR project using getFragmentsFromProject. Prior to fragment extraction, cell types in the human dataset were randomly downsampled to a maximum of 10,000 cells to reduce the computational cost. Per-cell-type peaks were called separately using MACS2 (--nomodel --shift -75 --extsize 150 --keep-dup all -q 0.01). The resulting per-cell-type peaks were intersected with the ArchR consensus peak set, retaining only those where the MACS2-derived summit directly overlapped an ArchR consensus peak. For each peak-cell-type combination, MACS2 q-values were converted into scores per million (SPM) values (Corces et al., 2018) by dividing the peak q-value by the sum of all q-values for that specific cell type and multiplying by 10⊠. Binary pseudobulk matrices were derived by applying an SPM threshold of ≥5. For human, we started from already processed snATAC-seq data (Orozco et al., 2023). Raw fragment files, corresponding cell metadata, and consensus peak calls were downloaded from Zenodo (7532115) and used to initialize an ArchR project. Per-cell-type peak calls and a binarized pseudobulk peak matrix were then generated as described for mouse. To expand astrocyte coverage, we complemented the matrix with two additional human brain astrocyte datasets (Corces et al., 2020; Morabito et al., 2021). Individual ArchR projects were initialized for each dataset by downloading the corresponding data (GEO:GSE147672 and Synapse:syn22130833). Astrocyte-specific peak calling was performed as previously described, converting MACS2 q-values to SPM values. We retained only peaks with SPM values ≥5 whose summits overlapped the human consensus retinal peak set, integrating them into the binarized pseudobulk matrix as separate columns.

### Synteny analysis of retinal peaks

Following specifications from ENCODE (Abascal et al., 2020), we performed reciprocal cross-species mapping of retinal chromatin accessibility peaks using the UCSC liftOver tool (Hinrichs et al., 2006) with a minimum ratio of bases that must remap (-minMatch) set to 0.5. To assess synteny across multiple resolutions, the analysis was performed at three levels: 1) the union of all accessible peaks across cell types (*i*.*e*., global), 2) per-cell-type accessible peaks, and 3) the approximately 2,500 DA peaks per cell type. For each level, human peaks were mapped to the mouse genome and intersected with mouse peaks using BEDTools intersect (version 2.31.1) (Quinlan & Hall, 2010) with the -wa - wb options, and vice versa. Reciprocal pairs were defined as cases where a human peak overlapped a mouse peak and the same mouse peak overlapped the original human peak.

### Fine-tuning Enformer models

We trained sequence-to-function models on the previously generated pseudobulk peak matrices using gReLU (version 1.0.4) (Lal et al., 2025). All models shared a base architecture derived from the pre-trained Enformer model (Avsec et al., 2021). Specifically, all transformer layers of the Enformer trunk except the first were removed, and output cropping was disabled. Model inputs consisted of 512-bp-long genomic sequences centered on the ArchR consensus peaks in the pseudobulk matrix. Peaks were restricted to autosomes, and any peaks overlapping blacklisted regions (Amemiya et al., 2019) were filtered out. The remaining peaks were split as follows: chromosome 10 was used for validation, chromosome 11 for the held-out test set, and all remaining autosomes were used for training. Models were trained for a maximum of 15 epochs using a batch size of 512 and the Adam optimizer (Kingma & Ba, 2014) with a learning rate of 1 × 10⊠⊠, and the loss function was set to binary cross-entropy with logits (BCEWithLogitsLoss). The best model checkpoint was selected based on the minimum validation loss.

### CATlas model

A pre-trained chromatin accessibility model trained on the CATlas dataset (Zhang et al., 2021), spanning 204 different human cell types and tissues, was downloaded from the Hugging Face model hub (Genentech/human-atac-catlas-model).

### Retinal cis-regulatory element design

We designed *cis*-regulatory elements (CREs) using the human binary model and the gReLU library. For each cell type (AC, astrocytes, BC, HC, MG, photoreceptors, RGC, and RPE), we evolved 10 strong (SPM ≥20) differentially accessible (DA) peaks and 10 randomly generated sequences to maximize a cell-type specificity score termed MinGap (Gosai et al., 2024), which is defined as the difference between the minimum predicted accessibility all on-target cell types (*e*.*g*., cones and rods for photoreceptors) and the maximum predicted accessibility across all off-target cell types. Two complementary design strategies were applied: 1) directed evolution, an iterative greedy algorithm that introduces single-nucleotide substitutions for a maximum of 50 iterations; and 2) Ledidi (version 1.2.2) (Schreiber et al., 2025), a gradient-based sequence optimization approach run for a maximum of 20,000 iterations. Moreover, two additional directed evolution experiments were conducted specifically on MG to dissect the importance of the LHX2 transcription factor (TF). In the LHX2 motif penalization experiment, the algorithm was configured to penalize the presence or creation of LHX2 motifs on either strand with a weight of -0.1. Conversely, in the LHX2 motif insertion experiment, directed evolution was executed in pattern-insertion mode, allowing only insertions of the LHX2 consensus sequence (“TAATTA”) (Folgueras et al., 2013).

### Latent space embedding and projection

To visualize the relationship between endogenous DA peaks and designed CREs in the learned representation space of the human binary model, we generated sequence embeddings using gReLU (embed_on_dataset). Specifically, we embedded 2,500 DA peaks from ACs, astrocytes, BCs, HCs, MG, photoreceptors, and RGCs, as well as the designed CREs for each of these cell types, and extracted the internal representation for each sequence from the model’s penultimate layer. The resulting high-dimensional embedding vectors were flattened and reduced to 50 principal components by applying PCA from the scikit-learn package, and subsequently projected into two dimensions using the UMAP Python library (version 0.5.12) (McInnes et al., 2018).

### TF-MoDISco motif discovery

To identify sequence motifs driving cell-type-specific chromatin accessibility predictions, we applied TF-MoDISco (Shrikumar et al., 2018), as implemented in modisco-lite (version 2.3.2), to approximately 2,500 DA peaks for each retinal cell type and each model. Specifically, we computed per-nucleotide contribution scores using gradient-based saliency with gradient correction using gReLU (version 1.0.6; run_modisco). Attribution scores were computed using the same MinGap cell-type specificity scores used for CRE design. TF-MoDISco subsequently aggregated these per-position scores into recurring sequence patterns, which were matched to a non-redundant set of TF motif clusters (see below). Finally, sequence logos were generated utilizing a trim threshold of 0.2 (representing 20% of the maximum contribution score).

### Non-redundant transcription factor motif clusters

To obtain a non-redundant set of TF motifs, we clustered the JASPAR 2024 vertebrates collection (Rauluseviciute et al., 2023) by adapting a previously described procedure (Vierstra et al., 2020). First, pairwise motif similarity was assessed by running TOMTOM, as implemented in tangermeme (version 0.4.0) (Schreiber, 2025a), in an all-vs-all mode. The resulting E-values were log⊠⊠-transformed and clipped between -2 and 10. Motifs were then grouped using complete-linkage hierarchical clustering on the correlation distance of this matrix using fastcluster (version 1.2.6) (Müllner, 2013) and SciPy (version 1.15.3) (Virtanen et al., 2019), applying a distance cutoff of 0.7. Motifs forming clusters of size one were retained as singletons. For each multi-member cluster, a consensus position probability matrix (PPM) was derived by identifying a seed motif (defined as the member with the lowest median E-value relative to all other cluster members). The remaining motifs were aligned to this seed using the offsets and strands reported by TOMTOM, and the aligned PPMs were subsequently averaged. Finally, low-information-content positions were trimmed from both edges of each consensus matrix using gReLU, following the specifications from JASPAR (Rauluseviciute et al., 2023). The resulting clusters were annotated with a structured identifier formatted as CXXX:NAME:DBD. Specifically, gene symbols were extracted from the cluster members and mapped to gene-family stems using a custom curated lookup table, with up to the two most represented stems forming the NAME field. DNA-binding domain (DBD) classifications were extracted from the CIS-BP database (Weirauch et al., 2014), and the most frequently occurring structural family within each cluster was assigned to the DBD field. Singletons were assigned sequential cluster numbers immediately following the last multi-member cluster.

### Attribution analysis of Müller glia elements

To assess the functional importance of LHX2 motifs in the starting and designed MG CREs, we computed *in silico* mutagenesis (ISM) attribution scores for each nucleotide position using gReLU. Predictions were transformed using the log⊠ fold-change of the MinGap specificity score for MG. LHX2 motifs were identified by scanning each sequence with FIMO, as implemented in memelite (version 0.2.0) (Schreiber, 2025b), against the non-redundant motif cluster containing LHX2 (C100:HOXB_LHX:Homeodomain; see below) at a p-value threshold of 10⊠3. Finally, ISM scores were visualized as sequence logos, with predicted LHX2 motif positions and point mutations introduced during directed evolution highlighted.

### AAV reporter construct design and production

The AAV transfer plasmid was designed with the backbone and AAV2 ITRs from pAAV_CBh_CRE.T2A.BFP (GenScript). Between the ITRs, we inserted: a minimal promoter (minP), EGFP, WPRE, SV40 poly(A) signal, sCAG promoter, mCherry, and bGH poly(A) signal. Restriction enzyme sites were engineered to enable subsequent cloning: XbaI and AgeI sites upstream of minP for CREs/enhancers, and an NheI site between minP and EGFP for promoter sequences. The empty AAV transfer plasmid (AAV2_eReporter_empty) was synthesized by GenScript. CREs were cloned upstream of minP by digesting the empty vector with XbaI and AgeI, followed by NEBuilder HiFi DNA Assembly (NEB) of gene fragments synthesized by Twist Bioscience. The hRGR promoter was cloned by digesting the empty vector with XbaI and NheI, replacing the minP with the hRGR gene fragment synthesized by Twist Bioscience. Resulting AAV reporter plasmids were sent to PackGene Biotech for research-grade AAV packaging and production (AAV2.7m8 serotype).

### Mouse in vivo intravitreal injections

To validate the rAAV constructs *in vivo*, we performed intravitreal injections (IVT) of rAAVs in mice (42 total, 13 groups, 3-4 animals per group). IVT was performed on both eyes of 9-10-week-old male C57Bl/6 mice (stock #00064, Jackson Laboratory) on day 0, with each assigned promoter/element in a rAAV vector. Mice were anesthetized with 75-80 mg/kg ketamine and 15 mg/kg xylazine IP in 150-300 ⊠L sterile saline. A single dose of Sustained Release buprenorphine (Ethiqa XR) was administered at 3.25mg/kg at the time of anesthesia and provided analgesia for the next 3 days. BETADINE 5% Sterile Ophthalmic Prep Solution (Alcon Laboratories, Inc.) was applied to the surface area 5 minutes before the surgery. Under a surgical microscope, a 33 gauge needle was used to punch the eye at the posterior limbus. A 1mm glass tip with a hand-held Hamilton syringe was then used to deliver 2 ⊠L of 1 × 10^10^ to 1 × 10^13^ genomic particles rAAV through the existing entrance site. Glass tips were switched between the groups. Neomycin and polymyxin B sulfate and bacitracin zinc ophthalmic ointment (Bausch & Lomb) was applied to the injected eye post-injection, and the mice were placed on a pre-warmed warming plate at 37°C until they were fully awakened.

### Mouse retina sample collection

Mouse eyes were harvested at day 36 post-injection, enucleated, and fixed in 4% paraformaldehyde (PFA) at room temperature for 2 hours. Following fixation, the eyes were washed in phosphate-buffered saline (PBS), dehydrated through a graded ethanol series, cleared in xylene, and embedded in paraffin. Retinal sections were cut at a thickness of 5 ⊠m using a microtome and mounted onto glass slides for subsequent histological analyses.

### Immunofluorescence and image analysis

Retinas were incubated for 1 hour in blocking solution containing 5% normal goat serum (NGS) and 0.3% Triton X-100 in PBS, followed by overnight incubation at 4°C with primary antibodies diluted in antibody solution containing 1% BSA and 0.3% Triton X-100 in PBS. Primary antibodies used were rabbit anti-SOX9 (Millipore, AB5535, 1:1000) and chicken anti-GFP (Aves Labs, GFP-1020, 1:500). Samples were washed three times in PBS for 10 minutes each and then incubated for 2 hours at room temperature with donkey anti-rabbit Alexa Fluor 647 and donkey anti-chicken Alexa Fluor 488 secondary antibodies diluted 1:500 in antibody solution (1% BSA, 0.3% Triton X-100 in PBS). Slides were subsequently washed three times in PBS for 10 minutes each, with DAPI (Thermo Scientific, 62248, 1:200) included in the final wash for nuclear staining, and mounted with glass coverslips for imaging.

Slides were imaged using an Olympus SLIDEVIEW™ VS200W whole slide scanner at 20x magnification, with images acquired in the DAPI, CFP, FITC, and Cy5 fluorescence channels. The resulting whole slide images (WSIs) were converted to the OME-Zarr format (Moore et al., 2023) for subsequent analysis and processing. A two-step analysis procedure was conducted at different magnifications. The first step employed low magnification (2.5x) imaging to segment the ganglion cell layer (GCL), the inner nuclear layer (INL), and the outer nuclear layer (ONL) using a hybrid approach, combining pathologist-defined annotations with a previously trained machine learning model trained to segment the INL and ONL. In the second step, full resolution (20x) images were analyzed using Cellpose-SAM (Pachitariu et al., 2025) retrained via human-in-the-loop (Pachitariu & Stringer, 2022) annotations to identify GFP+ and Sox9+ regions within the GCL, INL, and ONL. An area-based segmentation approach was used to quantify co-localization without ambiguity from cell-boundary definitions. CRE specificity was defined as the percentage of GFP+ area that was Sox9+, and transduction efficiency as the percentage of Sox9+ area that was GFP+. Throughout the analysis, various binary morphological filters and small pixel area removals were applied using scikit-image (Walt et al., 2014). Due to memory constraints, most of the full resolution analysis was performed using a patch-based sliding window with overlap using Dask (Team, 2016).

## Code and Data Availability

The mouse multiome data generated in this study has been deposited in the GEO database under accession number GSE338378. The processed RNA and ATAC components, including the gene count matrix, the lineage-specific scATAC-seq peak matrix, cell type annotations, as well as the set of non-redundant transcription factor motifs are available via Zenodo (21270365). The code used in this work to process the multiome data, train models, and evolve and interpret elements, alongside the evolved elements themselves, is also available via Zenodo. The retinal models and corresponding datasets are available through the Hugging Face model hub (Genentech/retinal-models) and datasets hub (datasets/Genentech/retinal-datasets), respectively.

## Acknowledgements

We thank the members of Genentech’s “Good Enough Genomics” and ReLU teams, especially Caitlin Simopoulos, Luli Zou, Surag Nair, Laura Gunsalus, and M. Hasan Çelik, as well as Catherine McKenzie, and Florian Noack from Roche for their advice and useful feedback.

## Ethics Statement

All animal experiments were approved by the Genentech IACUC and conducted in compliance with the Institute for Laboratory Animal Research (ILAR) guidelines for the humane care and use of laboratory animals, and adhered to the Association for Research in Vision and Ophthalmology (ARVO) Statement for the Use of Animals in Ophthalmic and Vision Research.

## Notes

https://www.ncbi.nlm.nih.gov/geo/query/acc.cgi?acc=GSE338378

https://doi.org/10.5281/zenodo.21270366

https://huggingface.co/Genentech/retinal-models

https://huggingface.co/datasets/Genentech/retinal-datasets

### Competing Interest Statement

All authors are, or were at the time this work was conducted, employees of Genentech, Inc. or F. Hoffmann-La Roche AG, and may hold Roche stock or stock options.

### Summary of Updates

The mouse multiome data have been expanded from 9,211 to 54,073 cells and analyzed by Carlo De Donno. As a consequence, Figs. 1, 2B, and 3A, Figs. S1 and S2, and Table S1 have been modified, and Carlo De Donno has been added to the author list. The designed elements, which were optimized using the human model, are unchanged. The interpretation of the results does not change.

## References

Abascal, F., Acosta, R., Addleman, N. J., Adrian, J., Afzal, V., Ai, R., Aken, B., Akiyama, J. A., Jammal, O. A., Amrhein, H., Anderson, S. M., Andrews, G. R., Antoshechkin, I., Ardlie, K. G., Armstrong, J., Astley, M., Banerjee, B., Barkal, A. A., Barnes, I. H. A., … Weng, Z. (2020). Expanded encyclopaedias of DNA elements in the human and mouse genomes. Nature, 583(7818), 699–710. 10.1038/s41586-020-2493-4

Almeida, B. P. de, Schaub, C., Pagani, M., Secchia, S., Furlong, E. E. M., & Stark, A. (2024). Targeted design of synthetic enhancers for selected tissues in the Drosophila embryo. Nature, 626(7997), 207–211. 10.1038/s41586-023-06905-9

Alstyne, M. V., Tattoli, I., Delestrée, N., Recinos, Y., Workman, E., Shihabuddin, L. S., Zhang, C., Mentis, G. Z., & Pellizzoni, L. (2021). Gain of toxic function by long-term AAV9-mediated SMN overexpression in the sensorimotor circuit. Nature Neuroscience, 24(7), 930–940. 10.1038/s41593-021-00827-3

Amemiya, H. M., Kundaje, A., & Boyle, A. P. (2019). The ENCODE Blacklist: Identification of Problematic Regions of the Genome. Scientific Reports, 9(1), 9354. 10.1038/s41598-019-45839-z

Avsec, Ž., Agarwal, V., Visentin, D., Ledsam, J. R., Grabska-Barwinska, A., Taylor, K. R., Assael, Y., Jumper, J., Kohli, P., & Kelley, D. R. (2021). Effective gene expression prediction from sequence by integrating long-range interactions. Nature Methods, 18(10), 1196–1203. 10.1038/s41592-021-01252-x

Avsec, Ž., Latysheva, N., Cheng, J., Novati, G., Taylor, K. R., Ward, T., Bycroft, C., Nicolaisen, L., Arvaniti, E., Pan, J., Thomas, R., Dutordoir, V., Perino, M., De, S., Karollus, A., Gayoso, A., Sargeant, T., Mottram, A., Wong, L. H., … Kohli, P. (2026). Advancing regulatory variant effect prediction with AlphaGenome. Nature, 649(8099), 1206–1218. 10.1038/s41586-025-10014-0

Bainbridge, J. W. B., Smith, A. J., Barker, S. S., Robbie, S., Henderson, R., Balaggan, K., Viswanathan, A., Holder, G. E., Stockman, A., Tyler, N., Simon, P.-J., Bhattacharya, S. S., Thrasher, A. J., Fitzke, F. W., Carter, B. J., Rubin, G. S., Moore, A. T., & Ali, R. R. (2008). Effect of Gene Therapy on Visual Function in Leber’s Congenital Amaurosis. New England Journal of Medicine, 358(21), 2231–2239. 10.1056/nejmoa0802268

Chen, S., Wang, Q.-L., Nie, Z., Sun, H., Lennon, G., Copeland, N. G., Gilbert, D. J., Jenkins, N. A., & Zack, D. J. (1997). Crx, a Novel Otx-like Paired-Homeodomain Protein, Binds to and Transactivates Photoreceptor Cell-Specific Genes. Neuron, 19(5), 1017–1030. 10.1016/s0896-6273(00)80394-3

Corces, M. R., Granja, J. M., Shams, S., Louie, B. H., Seoane, J. A., Zhou, W., Silva, T. C., Groeneveld, C., Wong, C. K., Cho, S. W., Satpathy, A. T., Mumbach, M. R., Hoadley, K. A., Robertson, A. G., Sheffield, N. C., Felau, I., Castro, M. A. A., Berman, B. P., Staudt, L. M., … Zhu, J. (2018). The chromatin accessibility landscape of primary human cancers. Science, 362(6413). 10.1126/science.aav1898

Corces, M. R., Shcherbina, A., Kundu, S., Gloudemans, M. J., Frésard, L., Granja, J. M., Louie, B. H., Eulalio, T., Shams, S., Bagdatli, S. T., Mumbach, M. R., Liu, B., Montine, K. S., Greenleaf, W. J., Kundaje, A., Montgomery, S. B., Chang, H. Y., & Montine, T. J. (2020). Single-cell epigenomic analyses implicate candidate causal variants at inherited risk loci for Alzheimer’s and Parkinson’s diseases. Nature Genetics, 52(11), 1158–1168. 10.1038/s41588-020-00721-x

Eraslan, G., Avsec, Ž., Gagneur, J., & Theis, F. J. (2019). Deep learning: new computational modelling techniques for genomics. Nature Reviews Genetics, 20(7), 389–403. 10.1038/s41576-019-0122-6

Feng, J., Liu, T., Qin, B., Zhang, Y., & Liu, X. S. (2012). Identifying ChIP-seq enrichment using MACS. Nature Protocols, 7(9), 1728–1740. 10.1038/nprot.2012.101

Folgueras, A. R., Guo, X., Pasolli, H. A., Stokes, N., Polak, L., Zheng, D., & Fuchs, E. (2013). Architectural Niche Organization by LHX2 Is Linked to Hair Follicle Stem Cell Function. Cell Stem Cell, 13(3), 314–327. 10.1016/j.stem.2013.06.018

Fornes, O., Av-Shalom, T. V., Korecki, A. J., Farkas, R. A., Arenillas, D. J., Mathelier, A., Simpson, E. M., & Wasserman, W. W. (2023). OnTarget: in silico design of MiniPromoters for targeted delivery of expression. Nucleic Acids Research, 51(W1), W379–W386. 10.1093/nar/gkad375

Foti, S. B., Samulski, R. J., & McCown, T. J. (2009). Delivering multiple gene products in the brain from a single adeno-associated virus vector. Gene Therapy, 16(11), 1314–1319. 10.1038/gt.2009.106

Furukawa, T., Morrow, E. M., & Cepko, C. L. (1997). Crx, a Novel otx-like Homeobox Gene, Shows Photoreceptor-Specific Expression and Regulates Photoreceptor Differentiation. Cell, 91(4), 531–541. 10.1016/s0092-8674(00)80439-0

Galvan, A., Choi, D., Korecki, A. J., Gomes, A. de M., Fornes, O., Tanimura, J., Farkas, R. A., Lam, S. L., Lu, G., Petkau, T. L., Yao, A., Wasserman, W. W., Leavitt, B. R., Simpson, E. M., & Smith, Y. (2026). MiniPromoters Ple384 (TH) and Ple388 (PITX3) for targeting midbrain dopaminergic neurons in mice and monkeys. Scientific Reports, 16(1), 9277. 10.1038/s41598-026-37466-2

Goldman, D. (2014). Müller glial cell reprogramming and retina regeneration. Nature Reviews Neuroscience, 15(7), 431–442. 10.1038/nrn3723

Gomes, A. de M., Petkau, T. L., Korecki, A. J., Fornes, O., Galvan, A., Lu, G., Hill, A. M., Lam, S. L., Yao, A., Farkas, R. A., Wasserman, W. W., Smith, Y., Simpson, E. M., & Leavitt, B. R. (2024). New MiniPromoter Ple389 (ADORA2A) drives selective expression in medium spiny neurons in mice and non-human primates. Scientific Reports, 14(1), 28194. 10.1038/s41598-024-79004-y

Gosai, S. J., Castro, R. I., Fuentes, N., Butts, J. C., Mouri, K., Alasoadura, M., Kales, S., Nguyen, T. T. L., Noche, R. R., Rao, A. S., Joy, M. T., Sabeti, P. C., Reilly, S. K., & Tewhey, R. (2024). Machine-guided design of cell-type-targeting cis-regulatory elements. Nature, 634(8036), 1211–1220. 10.1038/s41586-024-08070-z

Granja, J. M., Corces, M. R., Pierce, S. E., Bagdatli, S. T., Choudhry, H., Chang, H. Y., & Greenleaf, W. J. (2021). ArchR is a scalable software package for integrative single-cell chromatin accessibility analysis. Nature Genetics, 53(3), 403–411. 10.1038/s41588-021-00790-6

Hao, Y., Stuart, T., Kowalski, M. H., Choudhary, S., Hoffman, P., Hartman, A., Srivastava, A., Molla, G., Madad, S., Fernandez-Granda, C., & Satija, R. (2024). Dictionary learning for integrative, multimodal and scalable single-cell analysis. Nature Biotechnology, 42(2), 293–304. 10.1038/s41587-023-01767-y

Hinrichs, A. S., Karolchik, D., Baertsch, R., Barber, G. P., Bejerano, G., Clawson, H., Diekhans, M., Furey, T. S., Harte, R. A., Hsu, F., Hillman-Jackson, J., Kuhn, R. M., Pedersen, J. S., Pohl, A., Raney, B. J., Rosenbloom, K. R., Siepel, A., Smith, K. E., Sugnet, C. W., … Kent, W. J. (2006). The UCSC Genome Browser Database: update 2006. Nucleic Acids Research, 34(suppl_1), D590–D598. 10.1093/nar/gkj144

Hoang, T., Wang, J., Boyd, P., Wang, F., Santiago, C., Jiang, L., Yoo, S., Lahne, M., Todd, L. J., Jia, M., Saez, C., Keuthan, C., Palazzo, I., Squires, N., Campbell, W. A., Rajaii, F., Parayil, T., Trinh, V., Kim, D. W., … Blackshaw, S. (2020). Gene regulatory networks controlling vertebrate retinal regeneration. Science, 370(6519), eabb8598. 10.1126/science.abb8598

Huber, W., Carey, V. J., Gentleman, R., Anders, S., Carlson, M., Carvalho, B. S., Bravo, H. C., Davis, S., Gatto, L., Girke, T., Gottardo, R., Hahne, F., Hansen, K. D., Irizarry, R. A., Lawrence, M., Love, M. I., MacDonald, J., Obenchain, V., Oleś, A. K., … Morgan, M. (2015). Orchestrating high-throughput genomic analysis with Bioconductor. Nature Methods, 12(2), 115–121. 10.1038/nmeth.3252

Hwang, D.-Y., Hwang, M. M., Kim, H.-S., & Kim, K.-S. (2005). Genetically engineered dopamine ⊠-hydroxylase gene promoters with better PHOX2-binding sites drive significantly enhanced transgene expression in a noradrenergic cell-specific manner. Molecular Therapy, 11(1), 132–141. 10.1016/j.ymthe.2004.08.017

Iyer, A., Fronteiro, R., Bhatia, P., Kumari, S., Singh, A., Zhou, J., Bocchi, R., Narayanan, R., & Tole, S. (2025). The transcription factor LHX2 suppresses astrocyte proliferation in the postnatal mammalian cerebral cortex. Development, 152(20). 10.1242/dev.204358

Jüttner, J., Szabo, A., Gross-Scherf, B., Morikawa, R. K., Rompani, S. B., Hantz, P., Szikra, T., Esposti, F., Cowan, C. S., Bharioke, A., Patino-Alvarez, C. P., Keles, Ö., Kusnyerik, A., Azoulay, T., Hartl, D., Krebs, A. R., Schübeler, D., Hajdu, R. I., Lukats, A., … Roska, B. (2019). Targeting neuronal and glial cell types with synthetic promoter AAVs in mice, non-human primates and humans. Nature Neuroscience, 22(8), 1345–1356. 10.1038/s41593-019-0431-2

Kingma, D. P., & Ba, J. (2014). Adam: A Method for Stochastic Optimization. arXiv. 10.48550/arxiv.1412.6980

Klimova, L., Antosova, B., Kuzelova, A., Strnad, H., & Kozmik, Z. (2015). Onecut1 and Onecut2 transcription factors operate downstream of Pax6 to regulate horizontal cell development. Developmental Biology, 402(1), 48–60. 10.1016/j.ydbio.2015.02.023

Kohn, D. B., Chen, Y. Y., & Spencer, M. J. (2023). Successes and challenges in clinical gene therapy. Gene Therapy, 30(10–11), 738–746. 10.1038/s41434-023-00390-5

Kwasnieski, J. C., Fiore, C., Chaudhari, H. G., & Cohen, B. A. (2014). High-throughput functional testing of ENCODE segmentation predictions. Genome Research, 24(10), 1595–1602. 10.1101/gr.173518.114

Lal, A., Garfield, D., Biancalani, T., & Eraslan, G. (2024). Designing realistic regulatory DNA with autoregressive language models. Genome Research, 34(9), 1411–1420. 10.1101/gr.279142.124

Lal, A., Gunsalus, L., Nair, S., Biancalani, T., & Eraslan, G. (2025). gReLU: a comprehensive framework for DNA sequence modeling and design. Nature Methods, 22(11), 2253–2257. 10.1038/s41592-025-02868-z

Li, C., & Samulski, R. J. (2020). Engineering adeno-associated virus vectors for gene therapy. Nature Reviews Genetics, 21(4), 255–272. 10.1038/s41576-019-0205-4

Li, J., Choi, J., Cheng, X., Ma, J., Pema, S., Sanes, J. R., Mardon, G., Frankfort, B. J., Tran, N. M., Li, Y., & Chen, R. (2024). Comprehensive single-cell atlas of the mouse retina. iScience, 27(6), 109916. 10.1016/j.isci.2024.109916

Li, J., Wang, J., Ibarra, I. L., Cheng, X., Luecken, M. D., Lu, J., Monavarfeshani, A., Yan, W., Zheng, Y., Zuo, Z., Colborn, S. L. Z., Cortez, B. S., Owen, L. A., Wick, B., Bao, X., Choi, J., Haeussler, M., Tran, N. M., Shekhar, K., … Chen, R. (2026). Single-cell atlas of the transcriptome and chromatin accessibility in the human retina. Nature Genetics, 58(2), 418–433. 10.1038/s41588-025-02454-1

Linder, J., Srivastava, D., Yuan, H., Agarwal, V., & Kelley, D. R. (2025). Predicting RNA-seq coverage from DNA sequence as a unifying model of gene regulation. Nature Genetics, 57(4), 949–961. 10.1038/s41588-024-02053-6

Lun, A. T. L., & Kancherla, J. (2023). Powering single-cell analyses in the browser with WebAssembly. Journal of Open Source Software, 8(89), 5603. 10.21105/joss.05603

Martin, J. F., & Poché, R. A. (2019). Awakening the regenerative potential of the mammalian retina. Development, 146(23), dev182642. 10.1242/dev.182642

McInnes, L., Healy, J., & Melville, J. (2018). UMAP: Uniform Manifold Approximation and Projection for Dimension Reduction. arXiv. 10.48550/arxiv.1802.03426

Melo, J. de, Zibetti, C., Clark, B. S., Hwang, W., Miranda-Angulo, A. L., Qian, J., & Blackshaw, S. (2016). Lhx2 Is an Essential Factor for Retinal Gliogenesis and Notch Signaling. The Journal of Neuroscience, 36(8), 2391–2405. 10.1523/jneurosci.3145-15.2016

Miesfeld, J. B., Gestri, G., Clark, B. S., Flinn, M. A., Poole, R. J., Bader, J. R., Besharse, J. C., Wilson, S. W., & Link, B. A. (2015). Yap and Taz regulate retinal pigment epithelial cell fate. Development, 142(17), 3021–3032. 10.1242/dev.119008

Moore, J., Basurto-Lozada, D., Besson, S., Bogovic, J., Bragantini, J., Brown, E. M., Burel, J.-M., Moreno, X. C., Medeiros, G. de, Diel, E. E., Gault, D., Ghosh, S. S., Gold, I., Halchenko, Y. O., Hartley, M., Horsfall, D., Keller, M. S., Kittisopikul, M., Kovacs, G., … Swedlow, J. R. (2023). OME-Zarr: a cloud-optimized bioimaging file format with international community support. Histochemistry and Cell Biology, 160(3), 223–251. 10.1007/s00418-023-02209-1

Morabito, S., Miyoshi, E., Michael, N., Shahin, S., Martini, A. C., Head, E., Silva, J., Leavy, K., Perez-Rosendahl, M., & Swarup, V. (2021). Single-nucleus chromatin accessibility and transcriptomic characterization of Alzheimer’s disease. Nature Genetics, 53(8), 1143–1155. 10.1038/s41588-021-00894-z

Müllner, D. (2013). fastcluster⊠: Fast Hierarchical, Agglomerative Clustering Routines for R and Python. Journal of Statistical Software, 53(9). 10.18637/jss.v053.i09

Novakovsky, G., Dexter, N., Libbrecht, M. W., Wasserman, W. W., & Mostafavi, S. (2023). Obtaining genetics insights from deep learning via explainable artificial intelligence. Nature Reviews Genetics, 24(2), 125–137. 10.1038/s41576-022-00532-2

Orozco, L. D., Owen, L. A., Hofmann, J., Stockwell, A. D., Tao, J., Haller, S., Mukundan, V. T., Clarke, C., Lund, J., Sridhar, A., Mayba, O., Barr, J. L., Zavala, R. A., Graves, E. C., Zhang, C., Husami, N., Finley, R., Au, E., Lillvis, J. H., … DeAngelis, M. M. (2023). A systems biology approach uncovers novel disease mechanisms in age-related macular degeneration. Cell Genomics, 3(6), 100302. 10.1016/j.xgen.2023.100302

Pachitariu, M., Rariden, M., & Stringer, C. (2025). Cellpose-SAM: superhuman generalization for cellular segmentation. bioRxiv, .04.28.651001. 10.1101/2025.04.28.651001

Pachitariu, M., & Stringer, C. (2022). Cellpose 2.0: how to train your own model. Nature Methods, 19(12), 1634–1641. 10.1038/s41592-022-01663-4

Park, K. S., Cho, Y. I., Mitragotri, S., & Zhao, Z. (2026). Viral vector⊠based gene therapies in the clinic: An update. Bioengineering & Translational Medicine, 11(1), e70106. 10.1002/btm2.70106

Quinlan, A. R., & Hall, I. M. (2010). BEDTools: a flexible suite of utilities for comparing genomic features. Bioinformatics, 26(6), 841–842. 10.1093/bioinformatics/btq033

Rauluseviciute, I., Riudavets-Puig, R., Blanc-Mathieu, R., Castro-Mondragon, J. A., Ferenc, K., Kumar, V., Lemma, R. B., Lucas, J., Chèneby, J., Baranasic, D., Khan, A., Fornes, O., Gundersen, S., Johansen, M., Hovig, E., Lenhard, B., Sandelin, A., Wasserman, W. W., Parcy, F., & Mathelier, A. (2023). JASPAR 2024: 20th anniversary of the open-access database of transcription factor binding profiles. Nucleic Acids Research, 52(D1), D174–D182. 10.1093/nar/gkad1059

Schreiber, J. (2025a). tangermeme: A toolkit for understanding cis-regulatory logic using deep learning models. bioRxiv, 2025.08.08.669296. 10.1101/2025.08.08.669296

Schreiber, J. (2025b). Tomtom-lite: accelerating Tomtom enables large-scale and real-time motif similarity scoring. Bioinformatics, 41(11), btaf577. 10.1093/bioinformatics/btaf577

Schreiber, J., Lorbeer, F. K., Heinzl, M., Reiter, F., Rafanel, B., Lu, Y. Y., Stark, A., & Noble, W. S. (2025). Programmatic design and editing of cis-regulatory elements. bioRxiv, 2025.04.22.650035. 10.1101/2025.04.22.650035

Shrikumar, A., Tian, K., Avsec, Ž., Shcherbina, A., Banerjee, A., Sharmin, M., Nair, S., & Kundaje, A. (2018). Technical Note on Transcription Factor Motif Discovery from Importance Scores (TF-MoDISco) version 0.5.6.5. arXiv. 10.48550/arxiv.1811.00416

Taskiran, I. I., Spanier, K. I., Dickmänken, H., Kempynck, N., Pančíková, A., Ekşi, E. C., Hulselmans, G., Ismail, J. N., Theunis, K., Vandepoel, R., Christiaens, V., Mauduit, D., & Aerts, S. (2024). Cell-type-directed design of synthetic enhancers. Nature, 626(7997), 212–220. 10.1038/s41586-023-06936-2

Team, D. D. (2016). Dask: Library for dynamic task scheduling. http://dask.pydata.org

Vierstra, J., Lazar, J., Sandstrom, R., Halow, J., Lee, K., Bates, D., Diegel, M., Dunn, D., Neri, F., Haugen, E., Rynes, E., Reynolds, A., Nelson, J., Johnson, A., Frerker, M., Buckley, M., Kaul, R., Meuleman, W., & Stamatoyannopoulos, J. A. (2020). Global reference mapping of human transcription factor footprints. Nature, 583(7818), 729–736. 10.1038/s41586-020-2528-x

Virtanen, P., Gommers, R., Oliphant, T. E., Haberland, M., Reddy, T., Cournapeau, D., Burovski, E., Peterson, P., Weckesser, W., Bright, J., Walt, S. J. van der, Brett, M., Wilson, J., Millman, K. J., Mayorov, N., Nelson, A. R. J., Jones, E., Kern, R., Larson, E., … Contributors, S. 1 0. (2019). SciPy 1.0--Fundamental Algorithms for Scientific Computing in Python. arXiv. 10.48550/arxiv.1907.10121

Walt, S. van der, Schönberger, J. L., Nunez-Iglesias, J., Boulogne, F., Warner, J. D., Yager, N., Gouillart, E., Yu, T., & contributors, scikit-image. (2014). scikit-image: image processing in Python. PeerJ, 2, e453. 10.7717/peerj.453

Wang, D., Tai, P. W. L., & Gao, G. (2019). Adeno-associated virus vector as a platform for gene therapy delivery. Nature Reviews Drug Discovery, 18(5), 358–378. 10.1038/s41573-019-0012-9

Wang, S. K., Deng, B., Nair, S., Ren, X., Li, J., Tijerina, J., Prakhar, P., Luo, Z., Nnebe, C., Kim, S. H., Zhou, Y., Shah, S. H., Davis, A., Mahajan, R., Qiao, Y., Zhou, Y., Zhang, J., Xue, Y., Goldberg, J. L., … Wang, S. (2026). Deep learning-guided design of cell type-specific AAV promoters. bioRxiv, 2026.01.13.699371. 10.64898/2026.01.13.699371

Weirauch, M. T., Yang, A., Albu, M., Cote, A. G., Montenegro-Montero, A., Drewe, P., Najafabadi, H. S., Lambert, S. A., Mann, I., Cook, K., Zheng, H., Goity, A., van Bakel, H., Lozano, J.-C., Galli, M., Lewsey, M. G., Huang, E., Mukherjee, T., Chen, X., … Hughes, T. R. (2014). Determination and Inference of Eukaryotic Transcription Factor Sequence Specificity. Cell, 158(6), 1431–1443. 10.1016/j.cell.2014.08.009

Wu, F., Li, R., Umino, Y., Kaczynski, T. J., Sapkota, D., Li, S., Xiang, M., Fliesler, S. J., Sherry, D. M., Gannon, M., Solessio, E., & Mu, X. (2013). Onecut1 Is Essential for Horizontal Cell Genesis and Retinal Integrity. The Journal of Neuroscience, 33(32), 13053–13065. 10.1523/jneurosci.0116-13.2013

Xiong, W., Wu, D. M., Xue, Y., Wang, S. K., Chung, M. J., Ji, X., Rana, P., Zhao, S. R., Mai, S., & Cepko, C. L. (2019). AAV cis-regulatory sequences are correlated with ocular toxicity. Proceedings of the National Academy of Sciences, 116(12), 5785–5794. 10.1073/pnas.1821000116

Zhang, K., Hocker, J. D., Miller, M., Hou, X., Chiou, J., Poirion, O. B., Qiu, Y., Li, Y. E., Gaulton, K. J., Wang, A., Preissl, S., & Ren, B. (2021). A single-cell atlas of chromatin accessibility in the human genome. Cell, 184(24), 5985–6001.e19. 10.1016/j.cell.2021.10.024

