## Supplementary material for "Deep learning design and *in vivo* validation of Müller glia-specific *cis*-regulatory elements": Note S1

### Supplementary Note 1: On the design of retinal *cis*-regulatory elements using regression models

We adapted our deep learning workflow for designing retinal *cis*-regulatory elements (CREs) to using regression models. Unlike the previous binary models that predict chromatin accessibility as “open” or “closed”, regression models predict accessibility signal, enabling a more fine-grained CRE optimization process with desired cell-type accessibility levels.

#### *Generating pseudobulk peak matrices for training regression models*

To derive pseudobulk matrices for training regression models, we relied on the same ArchR projects (version 1.0.3.1) [(Granja et al., 2021)](https://app.readcube.com/library/e1a99ebd-898f-41a9-b7a8-e47888f0c939/all?uuid=7933513318561706&item_ids=e1a99ebd-898f-41a9-b7a8-e47888f0c939:9e570892-1495-4fb0-9e47-48d4975aa04b) used for the binarized pseudobulk peak matrices. We aggregated per-cell-type raw pseudobulk peak counts (getGroupSE) and subsequently scaled them to a continuous range between 0 and 1 using the MinMaxScaler function from scikit-learn (version 1.7.2) [(Pedregosa et al., 2012)](https://app.readcube.com/library/e1a99ebd-898f-41a9-b7a8-e47888f0c939/all?uuid=6953516028324546&item_ids=e1a99ebd-898f-41a9-b7a8-e47888f0c939:85d7c8cf-8de9-44a8-8b6e-5f774f5a27c7).

#### *Fine-tuning regression models*

Species-specific regression models were trained on these scaled pseudobulk matrices using gReLU (version 1.0.4) [(Lal et al., 2025)](https://app.readcube.com/library/e1a99ebd-898f-41a9-b7a8-e47888f0c939/all?uuid=8501195820430623&item_ids=e1a99ebd-898f-41a9-b7a8-e47888f0c939:3a3c84cc-1800-4ccc-ac15-dd0603e2dd01), setting the loss function to mean squared error, and retaining the remaining specifications used for training the binary models.

#### *Designing CREs* *using regression models*

CREs were designed using the same complementary design strategies (directed evolution and Ledidi [(Schreiber et al., 2025)](https://app.readcube.com/library/e1a99ebd-898f-41a9-b7a8-e47888f0c939/all?uuid=6284744123535738&item_ids=e1a99ebd-898f-41a9-b7a8-e47888f0c939:3a2046b5-6a40-4475-bea5-ec1b2acb2d40)) and starting sequences per cell type. However, instead of the MinGap cell-type specificity score [(Gosai et al., 2024)](https://app.readcube.com/library/e1a99ebd-898f-41a9-b7a8-e47888f0c939/all?uuid=7156033594217867&item_ids=e1a99ebd-898f-41a9-b7a8-e47888f0c939:313b20e5-88b4-40d5-86db-fb7be20cbd1a), we used a ratio-based objective function that maximized the division between the predicted on-target and off-target accessibilities.

### References

[Gosai, S. J., Castro, R. I., Fuentes, N., Butts, J. C., Mouri, K., Alasoadura, M., Kales, S., Nguyen, T. T. L., Noche, R. R., Rao, A. S., Joy, M. T., Sabeti, P. C., Reilly, S. K., & Tewhey, R. (2024). Machine-guided design of cell-type-targeting cis-regulatory elements. *Nature*, *634*(8036), 1211–1220. https://doi.org/10.1038/s41586-024-08070-z
Granja, J. M., Corces, M. R., Pierce, S. E., Bagdatli, S. T., Choudhry, H., Chang, H. Y., & Greenleaf, W. J. (2021). ArchR is a scalable software package for integrative single-cell chromatin accessibility analysis. *Nature Genetics*, *53*(3), 403–411. https://doi.org/10.1038/s41588-021-00790-6
Lal, A., Gunsalus, L., Nair, S., Biancalani, T., & Eraslan, G. (2025). gReLU: a comprehensive framework for DNA sequence modeling and design. *Nature Methods*, *22*(11), 2253–2257. https://doi.org/10.1038/s41592-025-02868-z
Pedregosa, F., Varoquaux, G., Gramfort, A., Michel, V., Thirion, B., Grisel, O., Blondel, M., Müller, A., Nothman, J., Louppe, G., Prettenhofer, P., Weiss, R., Dubourg, V., Vanderplas, J., Passos, A., Cournapeau, D., Brucher, M., Perrot, M., & Duchesnay, É. (2012). Scikit-learn: Machine Learning in Python. *arXiv*. https://doi.org/10.48550/arxiv.1201.0490
Schreiber, J., Lorbeer, F. K., Heinzl, M., Reiter, F., Rafanel, B., Lu, Y. Y., Stark, A., & Noble, W. S. (2025). Programmatic design and editing of cis-regulatory elements. *bioRxiv*, 2025.04.22.650035. https://doi.org/10.1101/2025.04.22.650035](https://app.readcube.com/library/?style=APA%20Style%207th%20edition+%7B%22language%22:%22en-US%22%7D)
