## Supplementary material for "Deep learning design and *in vivo* validation of Müller glia-specific *cis*-regulatory elements": Figs. S1-S7

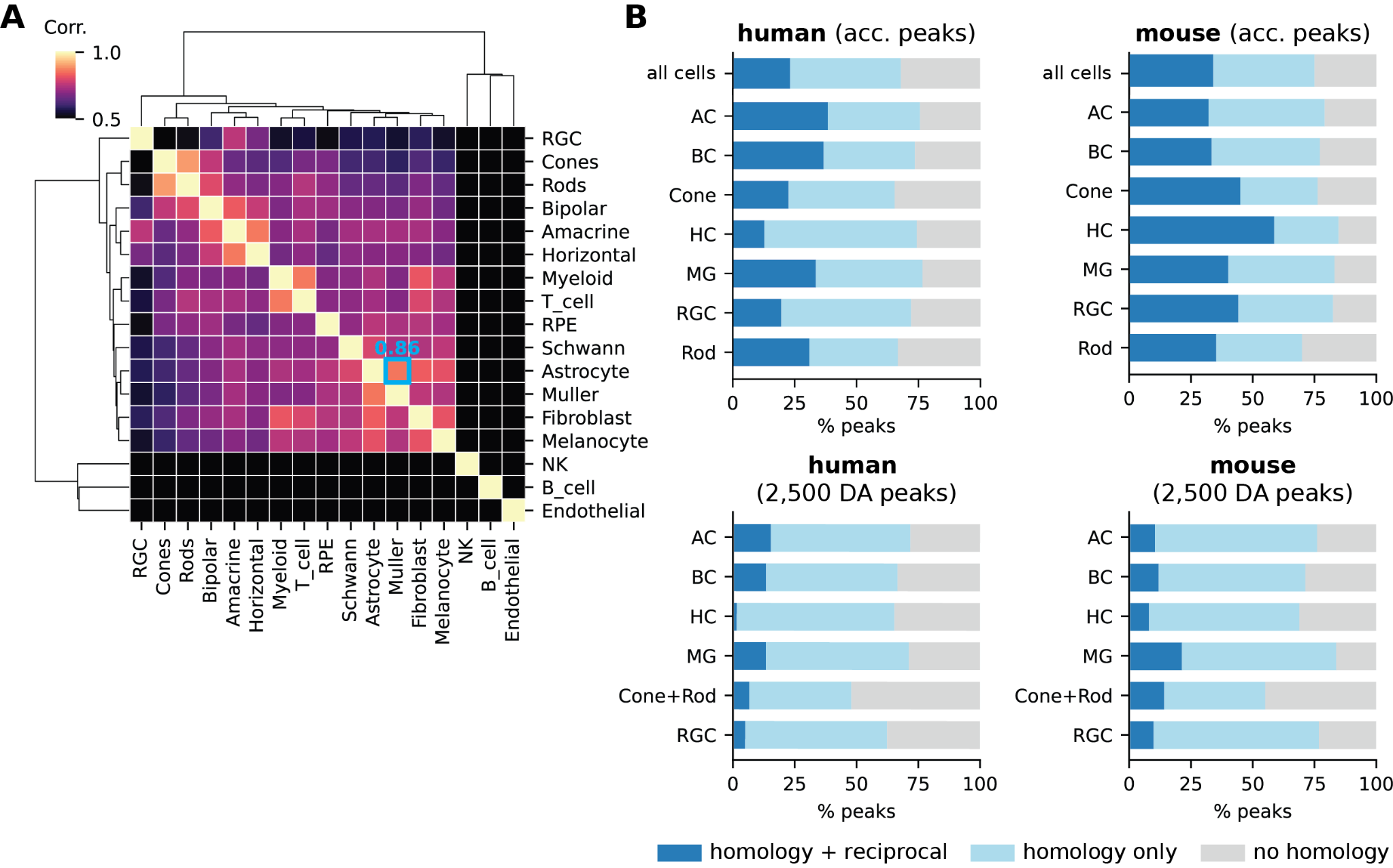

### *Figure S1*

(**A**) **Correlation of chromatin accessibility across human retinal cell types**.Hierarchical clustering of a cell-versus-cell Pearson correlation matrix based on binarized pseudobulk accessibility profiles. The high correlation between Müller glia and astrocytes (Corr. = 0.86) is highlighted in cyan. (**B**) **Synteny analysis of human and mouse retinal peaks.** Stacked bars show the conservation of peaks between human and mouse across three levels: 1) the union of all accessible peaks (all cells), 2) per-cell-type accessible peaks, and 3) approximately 2,500 differentially accessible (DA) peaks per cell type. Bars indicate the percentage of peaks with reciprocal homology (overlapping peaks identified in both species and in the same cells via bidirectional liftOver with minMatch = 0.5 [(Hinrichs et al., 2006)](https://app.readcube.com/library/e1a99ebd-898f-41a9-b7a8-e47888f0c939/all?uuid=2747770449108149&item_ids=e1a99ebd-898f-41a9-b7a8-e47888f0c939:14e5899d-cc2f-44bc-a255-7fe7a848b0e1); dark blue), non-reciprocal homology (one-way liftOver mapping; light blue), or no homology (gray).

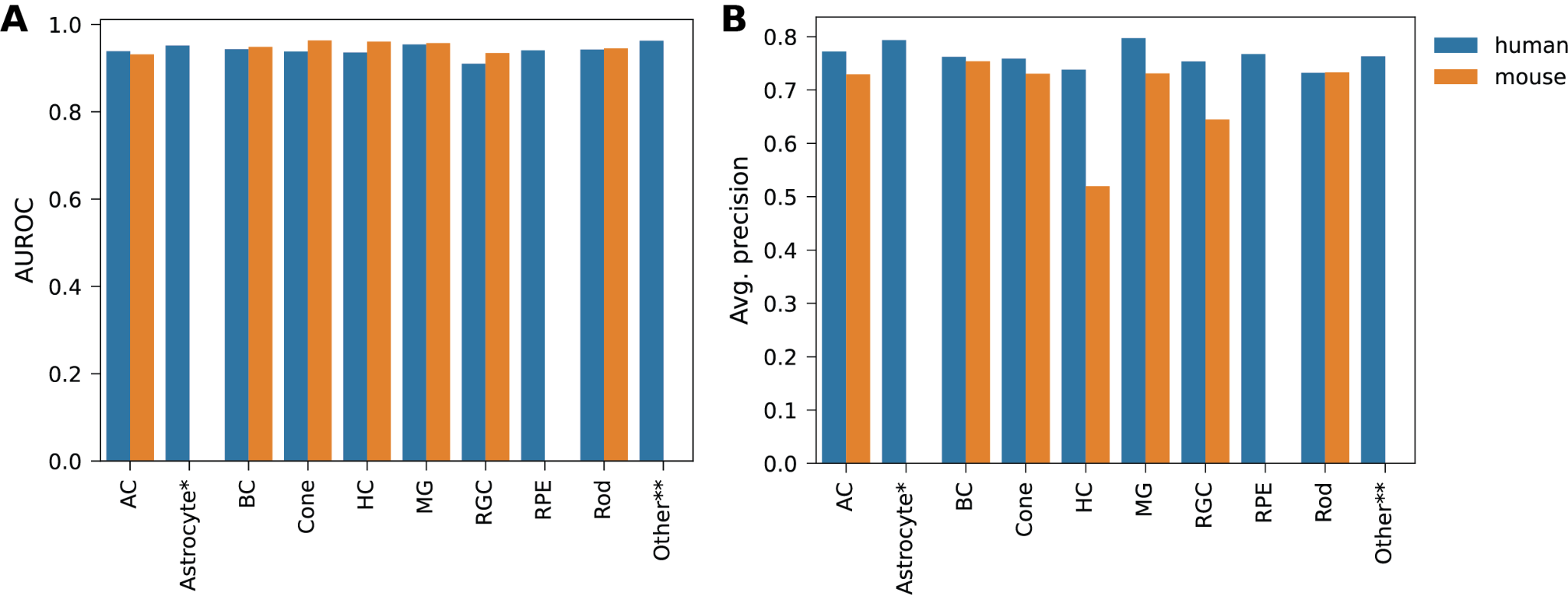

### *Figure S2*

Area Under the Receiver Operating Characteristic (AUROC; **A**) and average precision (**B**) computed on held-out chromosomes for the human (blue) and mouse (orange) models. AC, amacrine cells; BC, bipolar cells; HC, horizontal cells; MG, Müller glia; RGC, retinal ganglion cells; RPE, retinal pigment epithelium. One asterisk (*) denotes the three human astrocyte peak sets (one retinal and two brain-derived); two asterisks (**) denote non-retinal peak sets (*e.g.*, B and T cells).

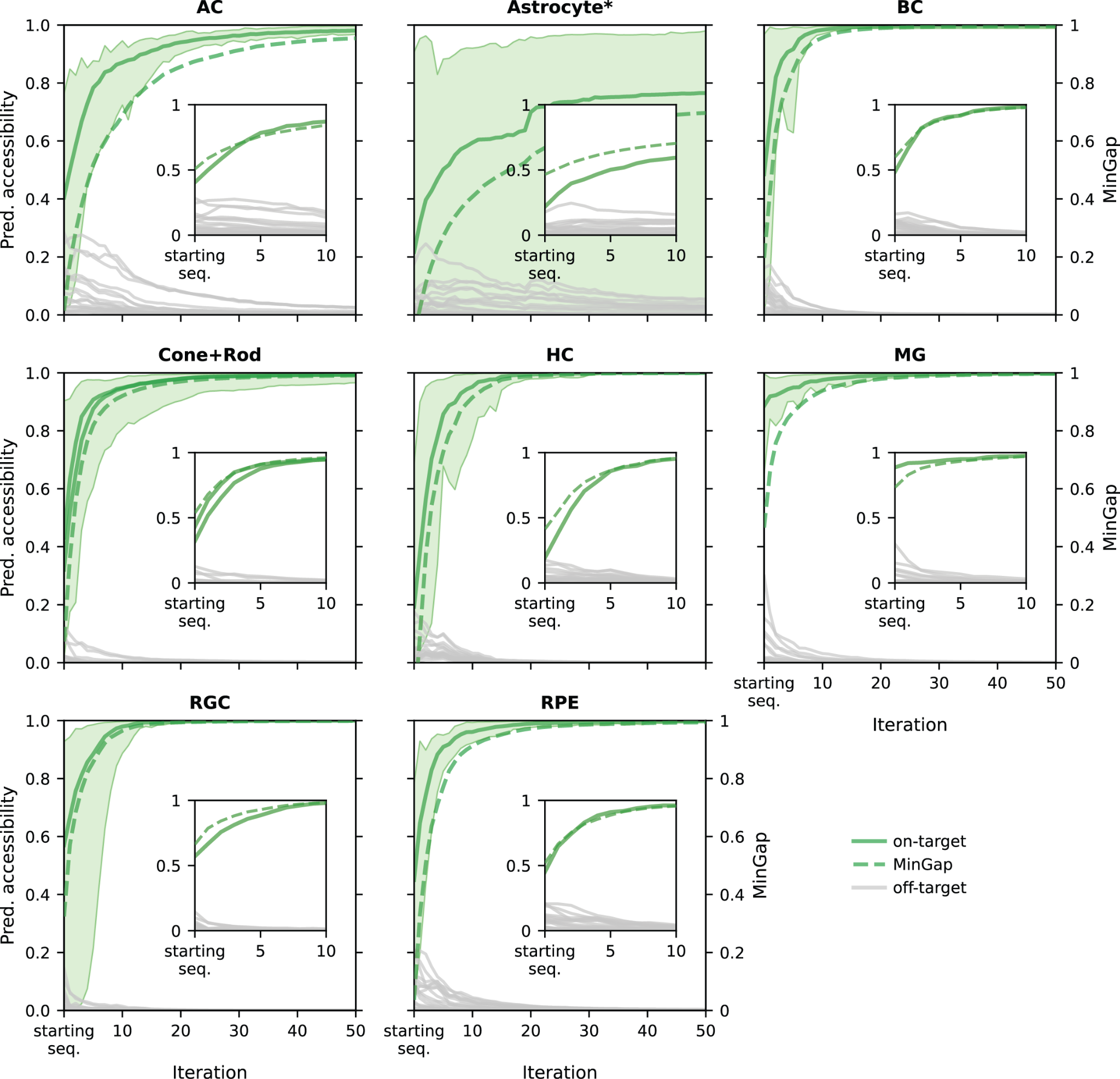

### *Figure S3*

Average predicted chromatin accessibility (shaded area indicates the min-max range) and MinGap score (secondary y-axis) in the human retinal model across 50 iterations of directed evolution and starting from endogenous DNA sequences that already partially exhibit the desired specificity. MinGap is defined as the difference between the minimum predicted on-target and the maximum predicted off-target accessibility in the human retina [(Gosai et al., 2024)](https://app.readcube.com/library/e1a99ebd-898f-41a9-b7a8-e47888f0c939/all?uuid=16880116103112064&item_ids=e1a99ebd-898f-41a9-b7a8-e47888f0c939:313b20e5-88b4-40d5-86db-fb7be20cbd1a). Insets highlights the rapid gain in predicted accessibility and specificity (*i.e.*, MinGap) within the first 10 iterations. AC, amacrine cells; BC, bipolar cells; HC, horizontal cells; MG, Müller glia; RGC, retinal ganglion cells; RPE, retinal pigment epithelium. One asterisk (*) denotes the three human astrocyte peak sets (one retinal and two brain-derived). Photoreceptors are denoted as “Cone+Rod”.

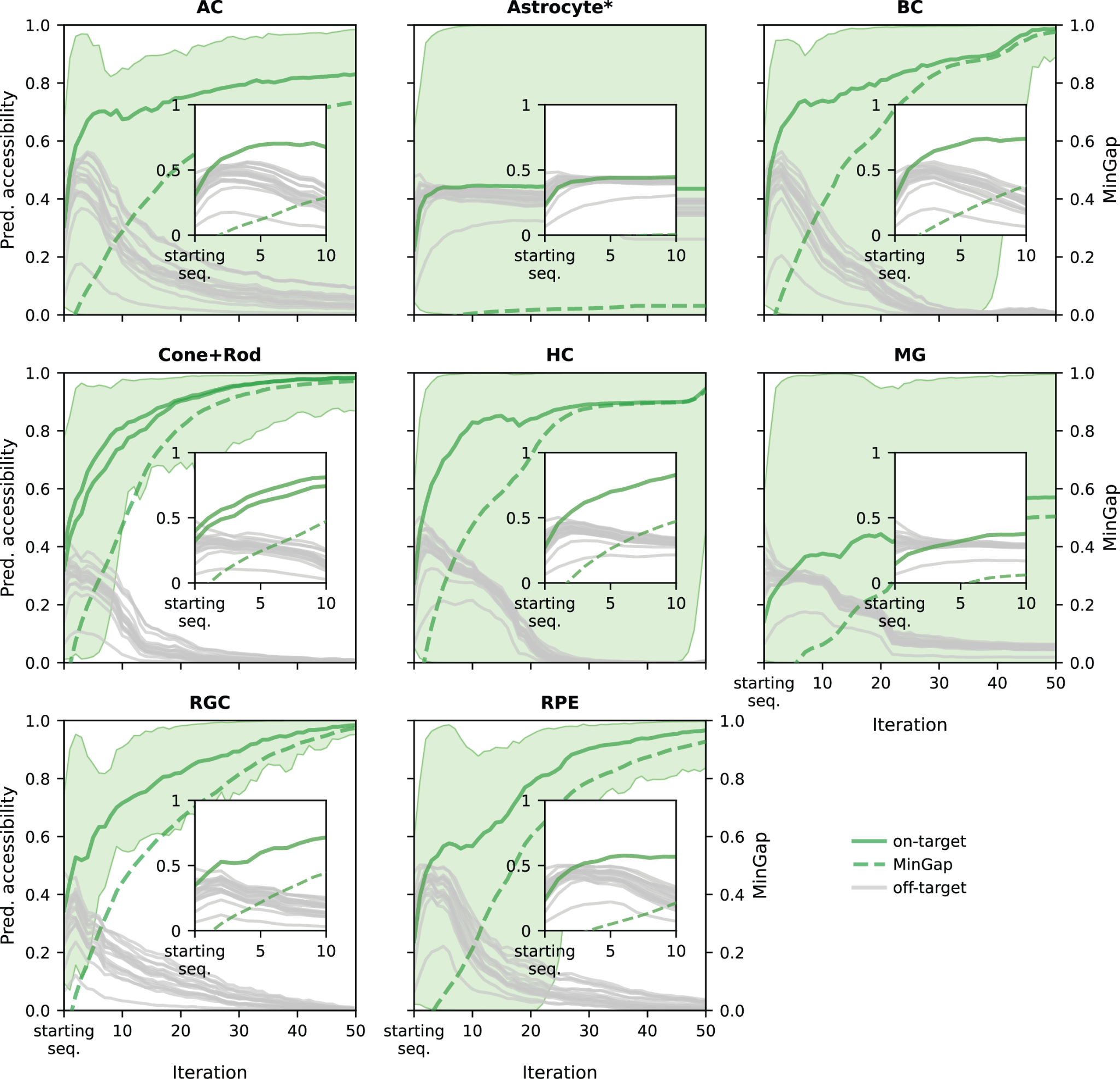

### *Figure S4*

Average predicted chromatin accessibility (shaded area indicates the min-max range) and MinGap score (secondary y-axis) in the human retinal model across 50 iterations of directed evolution and starting from random DNA. MinGap is defined as the difference between the minimum predicted on-target and the maximum predicted off-target accessibility in the human retina [(Gosai et al., 2024)](https://app.readcube.com/library/e1a99ebd-898f-41a9-b7a8-e47888f0c939/all?uuid=6599992233477722&item_ids=e1a99ebd-898f-41a9-b7a8-e47888f0c939:313b20e5-88b4-40d5-86db-fb7be20cbd1a). Insets highlights the rapid gain in predicted accessibility and specificity (*i.e.*, MinGap) within the first 10 iterations. AC, amacrine cells; BC, bipolar cells; HC, horizontal cells; MG, Müller glia; RGC, retinal ganglion cells; RPE, retinal pigment epithelium. One asterisk (*) denotes the three human astrocyte peak sets (one retinal and two brain-derived). Photoreceptors are denoted as “Cone+Rod”.

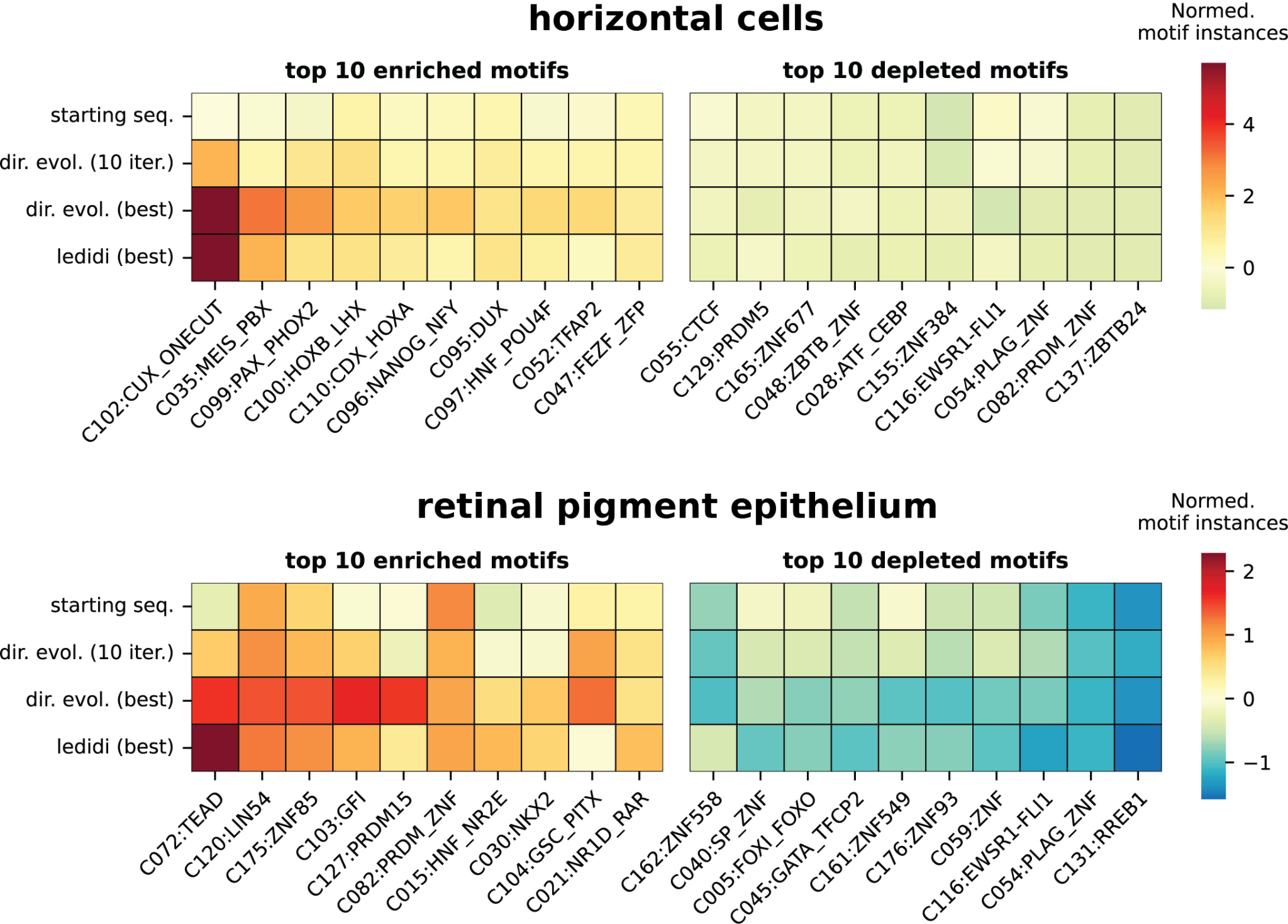

### *Figure S5*

Heatmaps showing the average normalized counts of the top 10 enriched (left) and depleted (right) transcription factor (TF) motifs in horizontal cells (top) and retinal pigment epithelium (bottom) for the starting sequences (starting seqs.) and best-evolved elements (*i.e.*, the highest-MinGap-scoring elements evolved from each starting sequence, where MinGap is defined as the difference between the minimum predicted on-target and maximum predicted off-target accessibility in the human retina [(Gosai et al., 2024)](https://app.readcube.com/library/e1a99ebd-898f-41a9-b7a8-e47888f0c939/all?uuid=4959174369893775&item_ids=e1a99ebd-898f-41a9-b7a8-e47888f0c939:313b20e5-88b4-40d5-86db-fb7be20cbd1a)) using directed evolution (dir. evol.) or Ledidi relative to approximately 2,500 differentially accessible peaks. For directed evolution, partially-evolved elements at iteration 10 (10 iter.) are also shown.

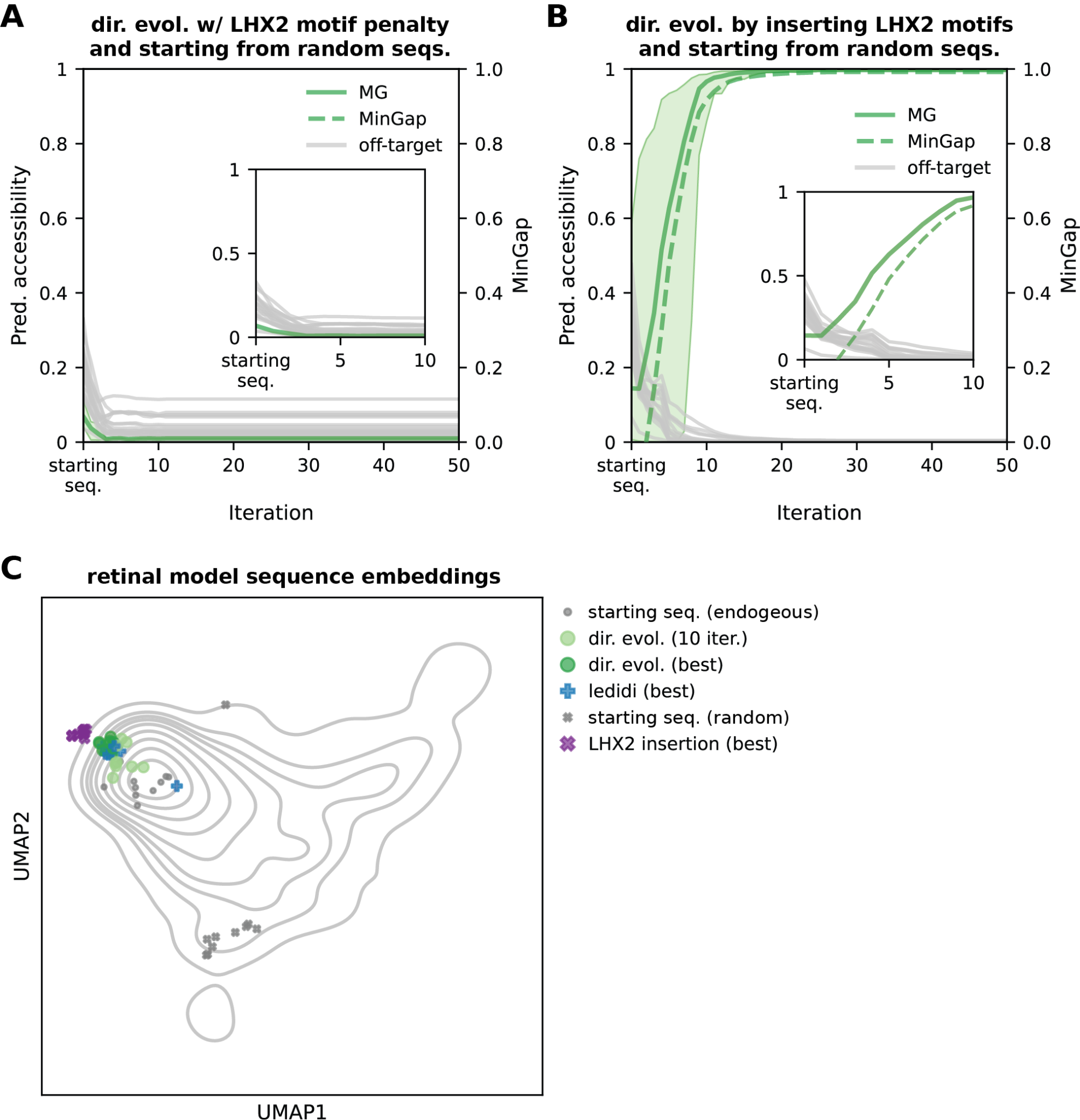

### *Figure S6*

**(A-B)** Average predicted chromatin accessibility (shaded area indicates the min-max range) and MinGap score (secondary y-axis) in the human retinal model across 50 iterations of directed evolution. MinGap is defined as the difference between the predicted Müller glia (MG) accessibility and the maximum predicted off-target accessibility in the human retina [(Gosai et al., 2024)](https://app.readcube.com/library/e1a99ebd-898f-41a9-b7a8-e47888f0c939/all?uuid=6757745924410841&item_ids=e1a99ebd-898f-41a9-b7a8-e47888f0c939:313b20e5-88b4-40d5-86db-fb7be20cbd1a). Directed evolution was performed by (**A**) starting from random DNA while penalizing the presence or creation of LHX2 motifs, or (**B**) allowing only insertions of the LHX2 consensus sequence (“TAATTA”) [(Folgueras et al., 2013)](https://app.readcube.com/library/e1a99ebd-898f-41a9-b7a8-e47888f0c939/all?uuid=286929653304586&item_ids=e1a99ebd-898f-41a9-b7a8-e47888f0c939:b6b9e256-9ecc-44f0-af7f-4cfce4e34348). Insets highlights the rapid gain in predicted accessibility and specificity (*i.e.*, MinGap) within the first 10 iterations. (**C**) UMAP projection of starting MG sequences (endogenous or random) and best-evolved elements (*i.e.*, the highest-MinGap-scoring elements evolved from each starting sequence) via directed evolution (dir. evol.), Ledidi, or LHX2 motif insertion into the MG differentially accessible peak landscape.

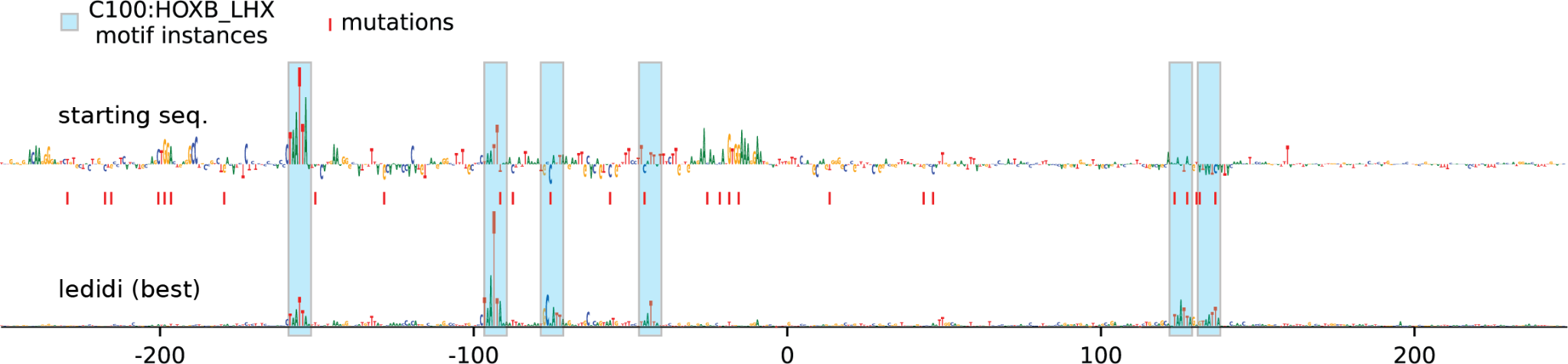

### *Figure S7*

Sequence optimization of a representative Müller glia (MG) element using Ledidi [(Schreiber et al., 2025)](https://app.readcube.com/library/e1a99ebd-898f-41a9-b7a8-e47888f0c939/all?uuid=4266499657304892&item_ids=e1a99ebd-898f-41a9-b7a8-e47888f0c939:3a2046b5-6a40-4475-bea5-ec1b2acb2d40). Attribution scores, representing the importance of each nucleotide for predicted MG accessibility, are shown as sequence logo heights. Red ticks along the x-axis indicate point mutations introduced relative to the starting sequence, and blue shaded regions denote identified C100:HOXB_LHX motif instances.
